# Alcohol reverses the effects of *KCNJ6* (GIRK2) noncoding variants on excitability of human glutamatergic neurons

**DOI:** 10.1101/2022.05.24.493086

**Authors:** Dina Popova, Isabel Gameiro-Ros, Mark M. Youssef, Petronio Zalamea, Ayeshia D. Morris, Iya Prytkova, Azadeh Jadali, Kelvin Y. Kwan, Chella Kamarajan, Jessica E. Salvatore, Xiaoling Xuei, David B. Chorlian, Bernice Porjesz, Samuel Kuperman, Danielle M. Dick, Alison Goate, Howard J. Edenberg, Jay A. Tischfield, Zhiping P. Pang, Paul A. Slesinger, Ronald P. Hart

## Abstract

Synonymous and noncoding single nucleotide polymorphisms (SNPs) in the *KCNJ6* gene, encoding G protein-gated inwardly rectifying potassium (GIRK2) channel subunit 2, have been linked with increased electroencephalographic frontal theta event-related oscillations (ERO) in subjects diagnosed with alcohol use disorder (AUD). To identify molecular and cellular mechanisms while retaining the appropriate genetic background, we generated induced excitatory glutamatergic neurons (iN) from iPSCs derived from four AUD-diagnosed subjects with *KCNJ6* variants (‘Affected: AF’) and four control subjects without variants (‘Unaffected: UN’). Neurons were analyzed for changes in gene expression, morphology, excitability and physiological properties. Single cell RNA sequencing suggests that *KCNJ6* AF variant neurons have altered patterns of synaptic transmission and cell projection morphogenesis. Results confirm that AF neurons express lower levels of GIRK2, have greater neurite area, and elevated excitability. Interestingly, exposure to intoxicating concentrations of ethanol induces GIRK2 expression and reverses functional effects in AF neurons. Ectopic overexpression of GIRK2 alone mimics the effect of ethanol to normalize induced excitability. We conclude that *KCNJ6* variants decrease GIRK2 expression and increase excitability and that this effect can be minimized or reduced with ethanol.

## Introduction

Alcohol use disorder (AUD) is a heritable (*h*^2^ = 0.49) condition characterized by an impaired ability to stop or control alcohol use despite adverse social, occupational, or health consequences^1^. An estimated 14.1 million American adults (ages 18+) were diagnosed with AUD in 2019 and the numbers continue to grow, particularly due to the COVID-19 pandemic^2^. Even though several evidence-based treatments are available for AUD^3–5^, there is a high discrepancy in treatment outcomes, suggesting the existence of a variety of traits influencing the development of specific physiological and behavioral responses to alcohol. Understanding biological factors, specifically genetic risk, at the level of neuronal function is fundamental to developing tailored preventive interventions and better matching of patients to treatment.

The Collaborative Study on the Genetics of Alcoholism (COGA) collected a diverse set of phenotypes, including electroencephalogram (EEG) parameters, in the search for genes and endophenotypes associated with AUD^6,7^. A family-based, genome-wide association study (GWAS) of a frontal theta event related oscillation (ERO) phenotype identified an association with several single nucleotide polymorphisms (SNPs), including a synonymous SNP, rs702859, in the *KCNJ6* gene on chromosome 21^8^. More recent studies concluded that this polymorphism influences the magnitude and topography of ERO theta power during reward processing in a monetary gambling task, reflecting a genetic link to neuronal circuits^9,10^. Two other SNPs (rs702860 and rs2835872) within *KCNJ6* were linked with the ERO endophenotype and AUD, but are noncoding, either intronic or within the 3’ untranslated region (3’ UTR), respectively. Additional studies identified various genetic loci correlating with selected EEG results but all consistently include alcohol dependence^11,12^. To understand how these SNPs might influence these phenotypes, it is essential to characterize the pathways and mechanisms by which genetic risk unfolds at molecular, cellular and network resolutions. Furthermore, since the *KCNJ6* SNPs are non-coding or synonymous, the optimal strategy would incorporate human genetic backgrounds of actual subjects to preserve potential effects of noncoding sequences.

*KCNJ6* encodes the G protein-gated inwardly rectifying potassium channel subunit 2 (GIRK2), which, when assembled into a GIRK channel, plays a role in regulating cell excitability^13^. The inward rectifying properties of GIRK are based on the ability to conduct potassium ions into the cell more easily than out, leading to a reduction in excitability^14,15^. GIRK function in neurons is associated with activation of G protein-coupled receptors (GPCRs), which leads to dissociation of Gβγ subunits from the G protein complex and binding to the channel^16^. In addition, this family of GIRK channels can be potentiated by ethanol concentrations (20-50 mM) relevant to human alcohol usage through direct interaction with a hydrophobic pocket on the channel^17,18^. Studies in model organisms also reveal the importance of GIRK function in AUD: mice lacking GIRK2 demonstrate reduced ethanol analgesia, greater ethanol-stimulated activity in open-field tests, increased self-administration of ethanol, and failure to develop a conditioned place preference for ethanol^19–21^. Thus, GIRK channels not only modulate the excitability of the neurons but also play an essential role in regulating responses to alcohol.

Recent studies have employed the use of subject-specific induced stem cell technology to model, characterize and elucidate mechanisms underlying various types of addiction disorders including AUD^22–25^. The generation of specific subtypes of cultured human neurons enables the study of formerly inaccessible functional properties using standard neuroscience methods^22,24–26^. The contribution of various *KCNJ6* SNPs was found to affect a variety of properties associated with neuronal function or sensitivity to drugs of abuse, suggesting targets for potential therapeutic interventions. Associating specific genetic variants with mechanisms contributing to addictive behaviors may produce more effective therapies tailored to individual genotypes.

We hypothesize that noncoding SNP variants in *KCNJ6* alter neuronal excitability, potentially contributing directly or indirectly to network-level endophenotypes such as ERO. Furthermore, since ethanol interacts directly with GIRK2, we predict that excitability will be modulated by ethanol. To test these hypotheses, we evaluated a human neuron model system that includes variant genotypes, so that specific intergenic or noncoding sequences as well as genetic backgrounds influencing these functions would be preserved. Therefore, we selected subjects from the NIAAA/COGA Sharing Repository, four with the ERO-associated allelic variant in *KCNJ6* and the presence of diagnosed alcohol dependence and four without the variant and unaffected for AUD and produced eight iPSC lines. iPSCs were reprogrammed into excitatory human neurons (iN) to investigate the potential contribution of the *KCNJ6* SNPs to AUD diagnosis and the ERO power endophenotype (Fig. 1A). We found that neurons from *KCNJ6* variant AUD-affected individuals demonstrated initial transcriptomic and morpho-physiological differences, mostly affecting excitability of the cells, which were paralleled by differences in GIRK2 expression levels. Ethanol exposure, conversely, induced GIRK2 expression, ameliorating differences in excitability. Moreover, by overexpressing *KCNJ6* we replicated effects of ethanol on neuronal excitability. The results promote a better understanding of the association between genetic variants and brain-wide changes affecting AUD risk and can potentially be used for development of personalized interventions.

**Figure 1.**
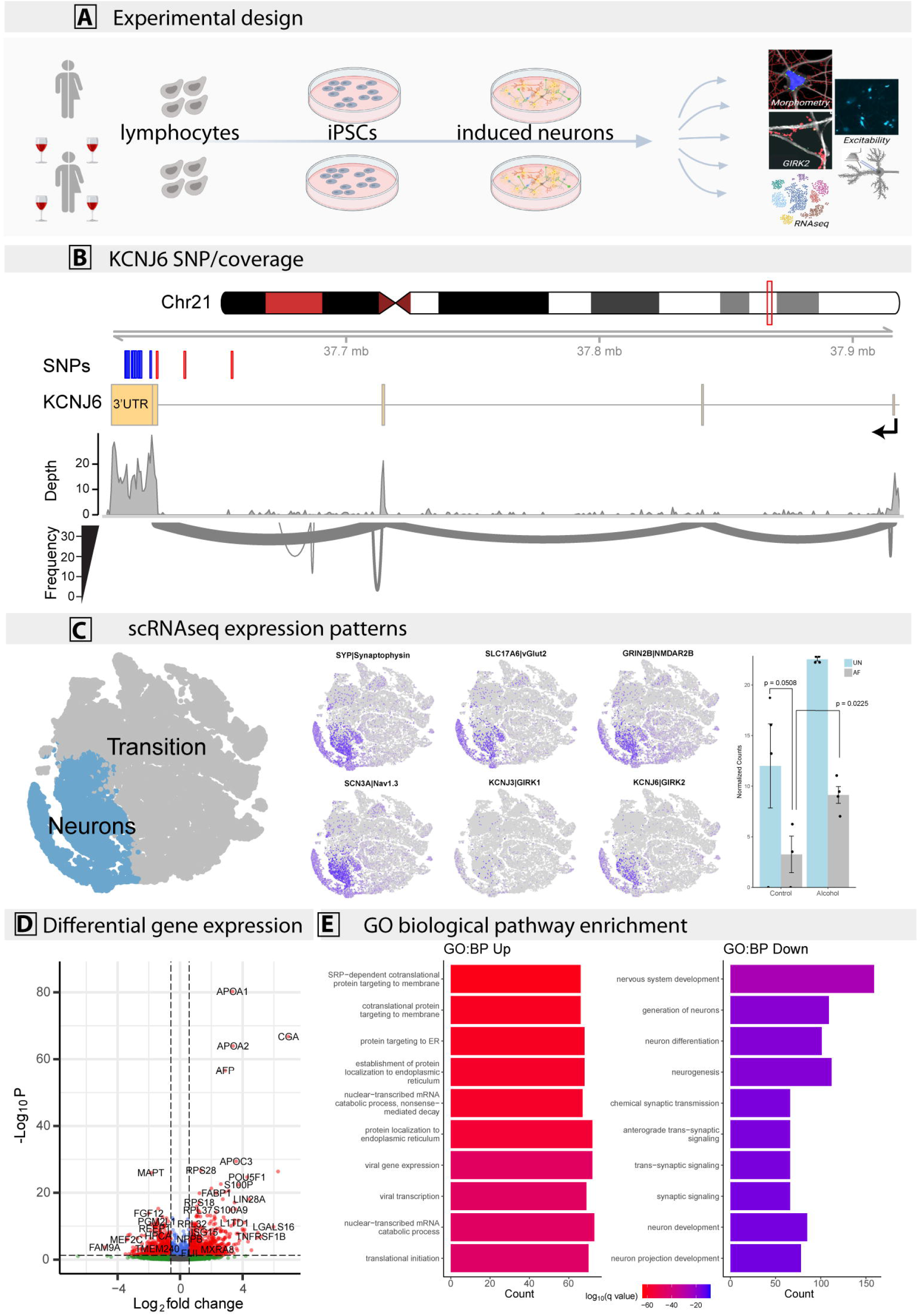
Experimental design and gene expression analysis. **A**. Diagram outlining experimental design. Lymphocytes from subjects with or without AUD diagnosis and *KCNJ6* haplotype variants were selected, reprogrammed into iPSC, induced into excitatory iNs, and analyzed by morphometry, immunocytochemistry, gene expression, and electrophysiology. **B**. Sequencing alignment and depth analysis of bulk RNA sequencing confirmed expression of *KCNJ6* mRNA in iN cultures, specifically the ENST00000609713 isoform, containing an 18.1 kilobase 3’UTR region. *KCNJ6* exons are mapped to chromosome locations (marked in mb, megabases) and the position of the gene is indicated by the red box on the chromosome 21 pictogram, top, with transcription direction on the minus strand indicated by the broken arrow. Variant analysis of RNA sequences predicts a region of linkage disequilibrium of 22 SNPs, including the 3 SNPs used to select subjects (red, Table 1), and 19 additional SNPs (blue, Supplemental Table 1). Depth: number of sequencing reads per base aligned by position. Frequency: thickness of curved lines represents the relative frequency of splice site utilization between exons. **C**. Single-cell RNAseq identifies a cluster of induced neurons (lower left), expressing markers consistent with neuronal function including *SYP, SLC17A6, GRIN2B, SCN3A, KCNJ3*, and *KCNJ6*; distinct from “transition neurons” that either do not express these markers or express sporadically. Additional markers are plotted in Supplemental Fig. 2E. Isolating *KCNJ6* mRNA expression, aggregated by subject and treatment, AF neurons expressed a trend towards lower levels than UN neurons (p = 0.0508; Wald test), but treatment of 7d with IEE at 20 mM peak concentration increased AF expression above untreated (p = 0.0225) to levels similar to UN control (p = 0.322, not denoted on figure). **D**. Volcano plot for untreated UN vs AF neurons, highlighting genes significantly different (FDR > 0.05) and at least 1.5-fold changed (red dots). Genes below the fold-change cut-off are marked in green, and those not significantly different are marked in blue. Significantly different genes are listed in Supplemental Table 3. **F**. Gene ontology (GO) enrichment of top 10 biological process (BP) terms for up- or down-regulated genes. Plots indicate the number of regulated genes from the term and the color indicates the adjusted p-value (q-value; key). All enriched terms are listed in Supplemental Tables 4-5.

## Methods

### Generation of human iPS cells and glutamatergic neurons (iN)

Subjects were selected from the NIAAA/COGA Sharing Repository of the Collaborative Study on the Genetics of Alcoholism (COGA) project with characteristics summarized in Table 1. Selection criteria consisted of *KCNJ6* SNP genotype, DSM IV diagnosis of AUD, sex, and availability of frozen lymphocytes in the repository. All subjects were of European ancestry. See **Supplemental Methods** for details. The previously-described protocol for generating glutamatergic human induced neurons (iN)^27–29^ was modified. To obtain cultures with enhanced levels of spontaneous activity, we minimized the time of Ngn2 induction to reduce alternate cell identities^30^ and fed cultures with a low-molality medium for at least 30 days post-induction^31^ as described in the **Supplemental Methods**. Cultures of iN typically exhibited resting membrane potentials averaging −40 mV with the presence of synaptic markers, spontaneous action potentials, and synaptic activity.

### Intermittent Ethanol Exposure (IEE)

Since ethanol evaporates during culture incubation (Supplemental Figure 7), we added ethanol to 20 mM and then replenished half the medium daily with 40 mM ethanol. Ethanol concentrations were measured using an AM1 Alcohol Analyzer (Analox Instruments, Ltd.)^25^.

### RNA sequencing

For bulk RNAseq, individual iN cultures were harvested in lysis buffer and column purified (Quick-RNA kit, Zymo). RNA was sent to Novogene, Inc., for sequencing. Analysis is described in the **Supplemental Methods**. RNAseq data have been deposited with the NIH GEO repository (accession number GSE196491). For single-cell RNA sequencing (scRNAseq), iN cultures were treated for 7 days using IEE^25^ starting 21 days after plating onto glia. At 28 days, iN were dissociated using trypLE Express (Thermo Fisher) for 5 minutes at 37°C and then processed and analyzed as described in the **Supplemental Methods**. These data have been deposited with the NIH GEO archive, accession number GSE203530.

### Electrophysiology

Analyses of iN used whole-cell patch-clamp electrophysiology as previously described^24,25^ with details provided in the **Supplemental Methods**. Drugs were applied through a perfusion system in the following concentrations: 160 nM ML297 or 20 mM ethanol. Clampfit software (pCIamp 11; Molecular Devices) was used for analysis of recordings.

### Immunocytochemistry and confocal imaging

iN cultures were fixed for 30 min in ice cold methanol and permeabilized using 0.2% Triton X-100 in PBS for 15 min at room temperature. Cells were then incubated in blocking buffer (5% BSA with 5% normal goat serum in PBS) for 30 min at room temperature and then incubated with primary antibodies diluted in blocking buffer overnight at 4°C, washed with PBS three times, and incubated with secondary antibodies for 1 h at room temperature. Confocal imaging was performed using a Zeiss LSM700. Primary antibodies used: rabbit anti-GIRK2 (Alomone labs, APC-006, 1:400), mouse anti-βIII-tub (BioLegend, MMS-435P,1:1000), chicken anti-MAP2 (Millipore AB5543, 1:1000), mouse anti-Syn1 (SYSY, 106-011, 1:200), mouse anti-PSD 95 (SYSY, 124-011,1:2000), mouse anti-mCherry (Thermofisher Scientific, M11217, 1:100).

### Fluorescent in situ hybridization (FISH)

Fluorescent, single molecule *in situ* hybridization was performed with the RNAscope® Multiplex Fluorescent Detection Kit v2 (Advanced Cell Diagnostics, ACD) following the manufacturer’s instructions and details provided in the **Supplemental Methods**.

### Image analysis

Cytometric analysis was performed using the Fiji/ImageJ image analysis program^32^ as described in the **Supplemental Methods**. ImageJ scripts can be downloaded from https://github.com/rhart604/imagej. The person performing analysis was blinded to genotype or line information.

### Lentiviruses

FSW-hSyn-GCaMP6f assembly was described previously^33^. FUGW-KCNJ6-mCherry was assembled using lentiviral backbone from FUGW (AddGene #14883), *KCNJ6* coding sequence amplified from human neuron cDNA (forward primer: agccaggaaaagcacaaaga, reverse primer: ggggagaagagaagggtttg), and mCherry with a nuclear localization signal from pME-nlsmCherry (a gift from Dr. Kelvin Kwan). During construction, the GIRK2 protein-coding sequence was tagged with a 3xHA tag and linked to mCherry with a T2A “self-cleaving” element.

### Calcium imaging in iPSC-derived iN populations

For assessing neuronal excitability, iN co-cultured with mouse glia on Matrigel™ coated 10 mm glass coverslips were transduced with lentivirus expressing hSyn-GCaMP6f at least 2 weeks prior to imaging to ensure robust expression. All fluorescence imaging experiments were performed in ^~^50 DIV (days in vitro) iNs. Details of the protocol and analysis are found in the **Supplemental Methods**.

### Statistics

Parameters that were sampled in cells from multiple microscopic fields and from multiple cell lines were fit to a linear mixed-effects model using the lme4 package in R^34^. The model included the cell line identifier and sex. A Tukey post-hoc test identified pairwise differences in two-factor models (e.g., genotype and ethanol). Electrophysiology results were modeled using generalized estimating equations (GEE) with the cell line as grouping identifier using the geepack R package^35^. Where appropriate, ANOVA or Student’s t-test was used as indicated in figure legends. For RNAseq, pseudo-bulk data (single cell reads pooled by subject identifier) were modeled in DESeq2^36^, testing first by likelihood ratio testing (LRT) over all groups, and then pairwise comparisons were evaluated using Wald tests. After Benjamini-Hochberg multiple measurements correction, a false discovery rate of 5% was set as a threshold for significance. Numbers of replicate cells and/or fields per cell line and genotype used for statistical testing are listed in Supplemental Table 6.

## Results

### Gene expression predicts functional differences between *KCNJ6* haplotypes

To investigate the role of *KCNJ6* gene variants in neuronal function and ethanol response we selected eight subjects of European ancestry from the COGA cell repository, with contrasting *KCNJ6* SNPs and the presence or absence of alcohol dependence (Table 1). Since multiple SNP genotypes were used, we label groups as AUD “affected” (AF) or “unaffected” (UF) for simplicity, since in this set of subjects diagnosis correlates with SNP haplotype. However, the focus of this study is the *KCNJ6* haplotype.

To confirm and extend SNP genotypes, iN cultures were harvested for bulk RNAseq analysis. To identify isoforms, RNAseq data identified only the longer *KCNJ6* isoform ENST00000609713 (Fig. 1B), which includes a 3’ exon of 18,112 nucleotides primarily consisting of 3’UTR. We identified 19 additional 3’UTR SNPs linked with the initial three (Supplementary Table 1; red lines in Fig. 1B), constituting a haplotype of 22 variants (synonymous and noncoding) within the expressed isoform of *KCNJ6* mRNA, all in a region of linkage disequilibrium (LD; Supplemental Figure 1C). There are no nonsynonymous variants, therefore, this haplotype is predicted to alter *KCNJ6* mRNA stability, translation, or other post-transcriptional processes.

Previous studies suggested that iN cultures may be heterogeneous, potentially confounding a bulk RNAseq analysis^30^. To enable analysis of iNs, we used single-cell RNAseq (scRNAseq) of pooled neurons from multiple cell lines, a strategy known as a “cell village^37^.” Based on preliminary results indicating differences in excitability, we cultured each haplotype separately to avoid potential secondary effects, combining cells each group in equivalent proportions. Following maturation (^~^30 days after induction), dissociated cells were processed to generate scRNAseq libraries. Mapping cells by t-distributed stochastic neighbor embedding (tSNE), a distinct cluster of cells coordinately expressed several markers consistent with neuron physiological function (Fig. 1C and Supplemental Fig. 2E), including synaptophysin (*SYP*), voltage-gated sodium channel (*SCN3A*), glutamate transporter (*SLC17A6*), and both NMDA (*GRIN2A, GRIN2B, GRIA2, GRIA4*) and kainate glutamate receptors (*GRIK2*). This cluster also expressed both *KCNJ6*, encoding GIRK2, and *KCNJ3*, encoding GIRK1, which are required to form heterotetrameric, functional channels^14,38^. Detection of mRNA in individual cells by scRNAseq underestimates expression, so the cluster identified as neurons likely expresses markers more uniformly than observed here. We conclude that this cluster of cells expresses components required for physiological activity, and that other cells, expressing subsets of these neuronal markers, are alternate products of induction, which we label as “transitional” neurons (Fig. 1C), since reprogramming is not identical to differentiation^39,40^. Therefore, we focused gene expression analysis on the phenotypically neuronal subset of cells.

Cells from individual subjects were distinguished by expressed SNPs^41^, and sequencing reads from each subject were combined to create a “pseudo-bulk” analysis, treating each subject and treatment condition as replicates. Comparing the untreated AF group with the untreated UN group, we identified 797 up-regulated genes, and 596 down-regulated genes (Fig. 1D; Supplemental Table 3). Examining the results for differences by sex instead of *KCNJ6* haplotype identified only 6 genes, 4 of which are encoded on X or Y, indicating that the sex of the samples did not substantially contribute to gene expression differences. Gene ontology analysis of the down-regulated genes (Fig. 1E; Supplemental Table 4-5) predicts several biological processes associated with nervous system development, axonal transport, and trans-synaptic signaling. By grouping enriched gene ontology terms by their parent terms (Supplemental Fig. 3), the major themes in the down-regulated genes (Supplemental Table 5) are synaptic signaling and neuro projection morphogenesis. Up-regulated genes (Supplemental Table 4) predict functions associated with protein targeting within the cells, catabolic metabolism, and nonsense-mediated decay. *KCNJ6* mRNA trended lower in untreated AF than in UN (Fig. 1C; p = 3.15 × 10^−5^ by LRT over both genotype and ethanol treatment; p = 0.0508 for untreated AF vs. UN by Wald test). Interestingly, 7 d of IEE increased levels of *KCNJ6* mRNA (Fig. 1C; p = 0.0225, Wald test) so that the treated AF group was no longer different from untreated UN (p = 0.322; Wald test). Analysis by sex identified no significant differences in *KCNJ6* mRNA. Results indicate that the variant *KCNJ6* haplotype leads to differential expression of GIRK2 and that ethanol exposure will reverse these effects.

### GIRK2 expression and function in iPSC-derived induced excitatory human neurons (iN)

Given the differences in gene expression and partial reversal of KCNJ6 downregulation, we evaluated detailed neuronal phenotypes in UN and AF iN cultures, including GIRK2 expression, neuronal morphology, and physiological response. Since GIRK2 immunocytochemistry had not been reported in cultured human neurons, we validated detection using mouse primary cortical cultures^42^ (Fig. 2A-B). Lentiviral-transduced GIRK2 overexpression increased levels of immunostaining about 3.5-fold (Fig. 2C, mCherry^+^ cells are KCNJ6 lentiviral-transduced). Importantly, silencing KCNJ6 with shRNAs or frameshift knockout by CRISPR/Cas9 eliminated GIRK2 immunoreactivity (Supplemental Fig. 4B,C), which further confirms antibody specificity.

**Figure 2.**
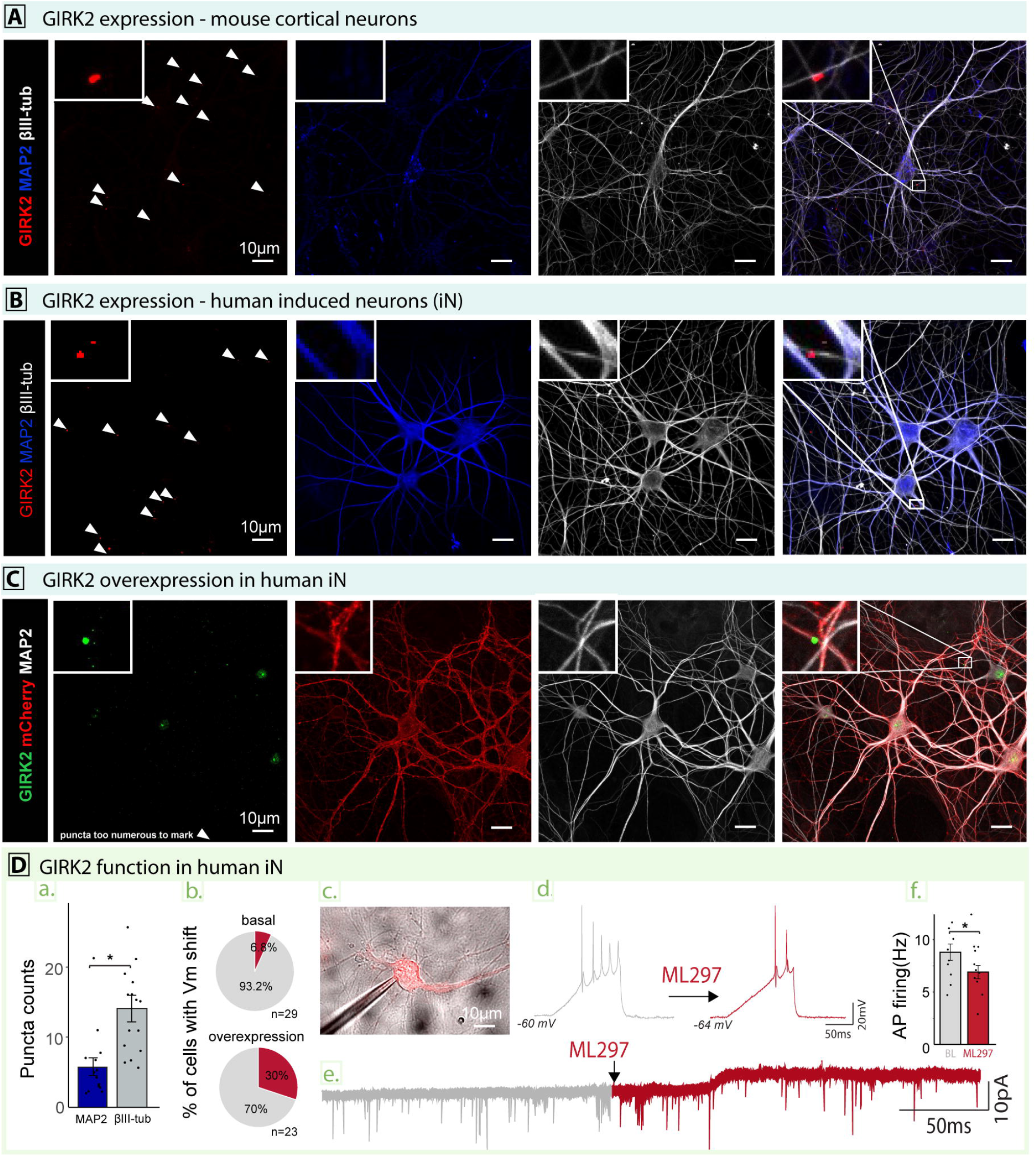
Validation of GIRK2 expression and function in human induced neurons. **A**. Representative confocal images of GIRK2 immunoreactivity in mouse cortical neurons. Arrowheads indicate locations of GIRK2-staining puncta, with an example punctum enlarged in the inset, overlapping or adjacent to βIII-tubulin-positive processes. We observed two cellular expression patterns – one where the entire neuron is decorated with GIRK2 antibody (Supplemental Fig. 4), or another where GIRK2 expression is relatively faint and observed mostly on neuronal processes, shown here. **B**. GIRK2 expression patterns in human induced neurons (iNs), showing representative confocal images from line 420. Inset shows two adjacent puncta. GIRK2 immunoreactivity matched a pattern of process-selective expression in mouse (Panel A and Supplemental Fig. 4A), where GIRK2 was detected as relatively small (^~^ 0.5 μm diameter) puncta scattered primarily along the processes. Localization of GIRK2 immunoreactivity in human iN did not directly colocalize with synaptic vesicle marker VGLUT2 or synaptic marker Syn1 (Supplemental Fig. 5), but instead was found most frequently adjacent to synapses but overlapping the shafts of the βIII tubulin-positive processes, and less so on MAP2 positive processes (Figs. 2D.a, 3C, 3E). Cultured neurons express βIII tubulin throughout the cell, but not as strongly in axonal processes^69^. We previously found that processes in human iN cells stained for ankyrin G, identifying the axonal initial segment, which similarly lacked βIII-tubulin^70^. Detection of GIRK2 primarily on βIII-tubulin^+^/MAP^-^ processes, therefore, suggests pre-axonal, and likely presynaptic, localization. **C**. Following infection of iN cultures with lentivirus expressing both *KCNJ6* and mCherry, large numbers of GIRK2^+^ puncta are seen in representative images (line 420). **D**. Evaluation of GIRK2 function in iNs, (a) quantification of GIRK2 expression on MAP2^+^ vs. βIII-tubulin^+^ neuronal processes. GIRK2 is more abundant on βIII-tubulin processes (p=0.0006, one-tailed Student’s t-test, n=15 cells per group, cell line 420). (b) Basal levels (upper pie plot) of the GIRK current in iNs as percent of neurons responding with hyperpolarization to the selective GIRK activator (160 nM ML297); compared with responding percentage when GIRK2 is overexpressed (lower pie chart). (c) Representative image of iN overexpressing GIRK2, as confirmed with mCherry fluorescence. (d) Representative traces of induced action potential firing before and after GIRK activation, demonstrating the contribution of GIRK function to cell excitability, (e) Representative trace of spontaneous postsynaptic potential (sEPSCs) recordings during ML297 (160 nM) GIRK activator wash-in, demonstrating a shift of 7mV holding current (amplifier-dependent compensation of GIRK-mediated membrane hyperpolarization). (f) Quantification of neuronal excitability at baseline and following GIRK activation with 160 nM ML297 (p=0.015, paired Student’s t-test, n=9 cells before/after ML297, cell line 376).

To confirm that GIRK2 expression contributes to potassium channel function, we treated iN cultures with ML297, a selective activator of the GIRK1/GIRK2 heterotetramer complex^43,44^. Results show that the magnitude and frequency of native (basal) currents observed in human iN were relatively small (Fig. 2D.b), with only 6.8% of the neurons responding to a shift in membrane potential holding current (^~^10 pA). However, cells overexpressing GIRK2 (identified by mCherry co-expression; Fig. 2D.c), exhibited an increased frequency of response (GIRK currents) to 30% of the cells after ML297 addition (Fig. 2D.b) without a change in magnitude (Fig. 2D.e). Importantly, GIRK activation affected excitability of the neurons by shifting resting membrane potential to more negative values (Fig. 2D.d), affecting the ability of neurons to fire APs when induced (Fig. 2D.f). We conclude that increased *KCNJ6* expression affects neuronal excitability by altering GIRK channel activity.

### *KCNJ6* haplotype alters morphology and membrane excitability

To identify neuronal properties affected by *KCNJ6* haplotype, we focused on three aspects: morphology, expression of GIRK2, and basal physiological properties. A detailed analysis of neuronal morphology is essential to rule out differences affecting excitability^45^. Most measures of basic neuronal morphology did not exhibit differences by haplotype, including soma size, circularity, and solidity, which describe the most fundamental aspects of neuron shape (Fig. 3B.a-c). However, neurite area, as determined by βIII-tubulin-staining, reflecting the number and/or branching of neurites per cell, increased in the AF group (p =0.018; Fig. 3B.d), with example images from individual cell lines in Figure 3C. Results are plotted for each cell line or aggregated by *KCNJ6* haplotype. As predicted by *KCNJ6* mRNA expression (Fig. 1J), GIRK2 immunoreactivity was decreased in the AF group (Fig. 3D), measured by puncta counts (p = 0.0012; Fig. 3D.a), puncta circularity (p =0.007; Fig. 3D.c), and solidity (p = 0.037; Fig. 3D.d) but not puncta size (Fig. 3D.b). No difference was found by sex. These results demonstrate an overall decrease in GIRK2 in neuronal processes in the AF (*KCNJ6* variant allele) group.

**Figure 3.**
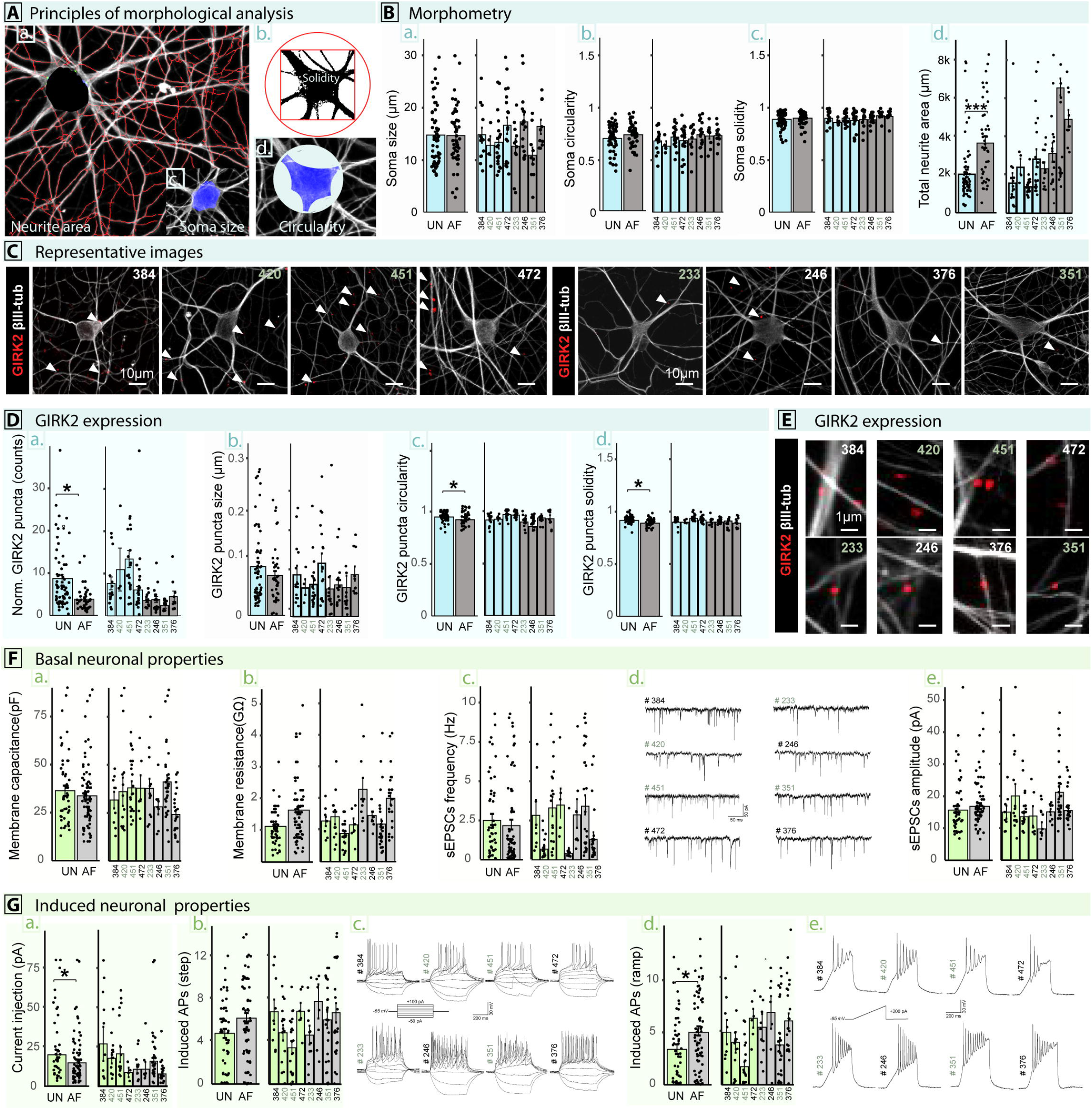
Impact of AUD-associated *KCNJ6* haplotype on neuronal properties. **A**. Principles of morphological analysis of induced neurons: (a) Neurite area was the total TuJ1^+^ (βIII-tubulin)^+^ staining area outside the cell soma. (b) Solidity is the area of the soma divided by its convex hull area. (c) Soma size was the area of the MAP2^+^ cell body. (d) Circularity compared the perimeter to the area. **B**. Morphometry of iNs from *KCNJ6* haplotype variant and affected (**AF**, cyan) or unaffected (**UN**, grey) individuals. Results are summed by group (left) or plotted individually by cell line (right), with subjects identified by line number (see Table 1—females identified with grey numbers). Individual cells are plotted as dots with the bar showing the mean, with error bars indicating the standard error of the mean (SEM). No significant differences were found in (a) soma size, (b) circularity, or (c) soma solidity, but total neurite area was increased in the AF group (p =0.018). **C**. Representative images of iNs from individual lines, with arrows identifying individual GIRK2 puncta (red) localized on βIII-tubulin^+^ processes (gray). **D**. GIRK2 expression was decreased in the AF as measured by puncta counts (a, p = 0.0012), circularity (c, p = 0.007), or solidity (d, p = 0.037), while there was no difference in puncta size (b). **E**. Representative images of individual GIRK2 puncta (red) localized on βIII-tubulin^+^ processes (gray). **F**. Electrophysiological analysis of passive neuronal properties, showing no difference in (a) membrane capacitance (b) membrane resistance, or (c) spontaneous EPSCs frequency. (d) Representative sEPSCs traces for each line, (e) Spontaneous EPSCs amplitude. **G**. Electrophysiological analysis of active neuronal properties. (a) Quantification of current required to shift resting membrane potential to −65mV in pA: difference by group p = 1.2 × 10^−5^; (b) quantification of maximum number of action potentials (APs) induced with the “step” protocol, p = 0.086; (c) representative traces of APs induced with the “step” protocol; (d) quantification of number of action potentials (APs) induced with the “ramp” protocol, p = 2.0 × 10^−9^; (e) representative traces of APs induced with the “ramp” protocol. A generalized linear model was used to evaluate group differences for morphometry and GIRK2 expression, and generalized estimation equations was used for electrophysiology results. Numbers of cell lines and replicates for each experiment are shown in Supplemental Table 6.

Reduced GIRK2 expression is predicted to affect the excitability of neurons. As expected, we found no differences in membrane capacitance, membrane resistance, spontaneous excitatory post-synaptic potential (sEPSC) frequencies or amplitudes, or spontaneous action potential (AP) firing (Fig. 3F). However, we observed increased excitability in the AF group (p = 1.2 × 10^−5^, Fig. 3G). Less current injection, using a ramp technique, was required for the AF group to shift the resting membrane potential (RMP) of neurons held at −65 mV, producing more AP firing, compared with the UN group (p = 2.0 × 10^−9^, Fig. 3G.d). Therefore, the *KCNJ6* variant haplotype group (AF) exhibited increased neurite area, reduced neurite GIRK2 expression, and greater excitability. Other neuronal properties were unchanged, suggesting that overall neuron differentiation status was similar.

Since cultures are heterogeneous (Fig. 1C), we wished to evaluate larger sample sizes. Therefore, we used calcium imaging to assess spontaneous and glutamate-stimulated firing. We selected four cell lines for these studies (AF: 233 and 246, and UN: 420 and 472) and used the viral-transduced, genetically-encoded calcium indicator GCaMP6f as a proxy for neuronal activity^46^. Example images show consistent expression in all cell lines (Fig. 4A). Cultures were treated with repeated pulses of 10 μM glutamate (each followed by wash-out with ACSF), and one pulse of 50 μM glutamate to elicit transient increases in fluorescence, indicating receptor-induced excitability, followed by pulses of 18 mM KCl, indicating cellular excitability (Fig. 4B-D). Neurons from both UN and AF groups had heterogeneous spiking patterns, with >50% of the neurons remaining inactive during the baseline period or after stimulation with glutamate or KCl. This is consistent with scRNAseq results indicating that only ^~^23% of the iN cells express genetic markers consistent with excitability (Fig. 1C). Increased spontaneous and induced spiking (> 1-2 spikes) was apparent in the AF group when viewing individual cell responses in a raster plot (Fig. 4B-C). Spiking frequencies increased 2.4-fold in baseline conditions (Fig. 4E; p = 1.1 × 10^−6^) and 1.3-fold following glutamate stimulation (Fig. 4F; p = 1.6 × 10^−3^) in the AF group, but no difference was found in KCl-induced activity (Fig. 4G; p = 0.85), confirming the above results from individual neurons. When examining the AF or UN neurons individually, we also observed higher spontaneous and glutamate-elicited excitability for lines 233 and 246 when compared to line 472, which was found to be the least active spontaneously or under glutamate stimulation among the four studied (Supp. Fig. 6). Overall, cells from the AF *KCNJ6* variant allele group were more excitable, with no difference in basal physiological properties (i.e., KCl-induced activity). Furthermore, not only did the spike rate increase per cell (bar plots, Spikes/ROI/min), but we also observed increases in the proportion of cells in a field that responded (pie plots). These results not only confirm findings of differences in excitability, but also demonstrate that the difference in the AF iNs is found in not only individual cell activity but also in the frequency of detecting active cells within a population.

**Figure 4.**
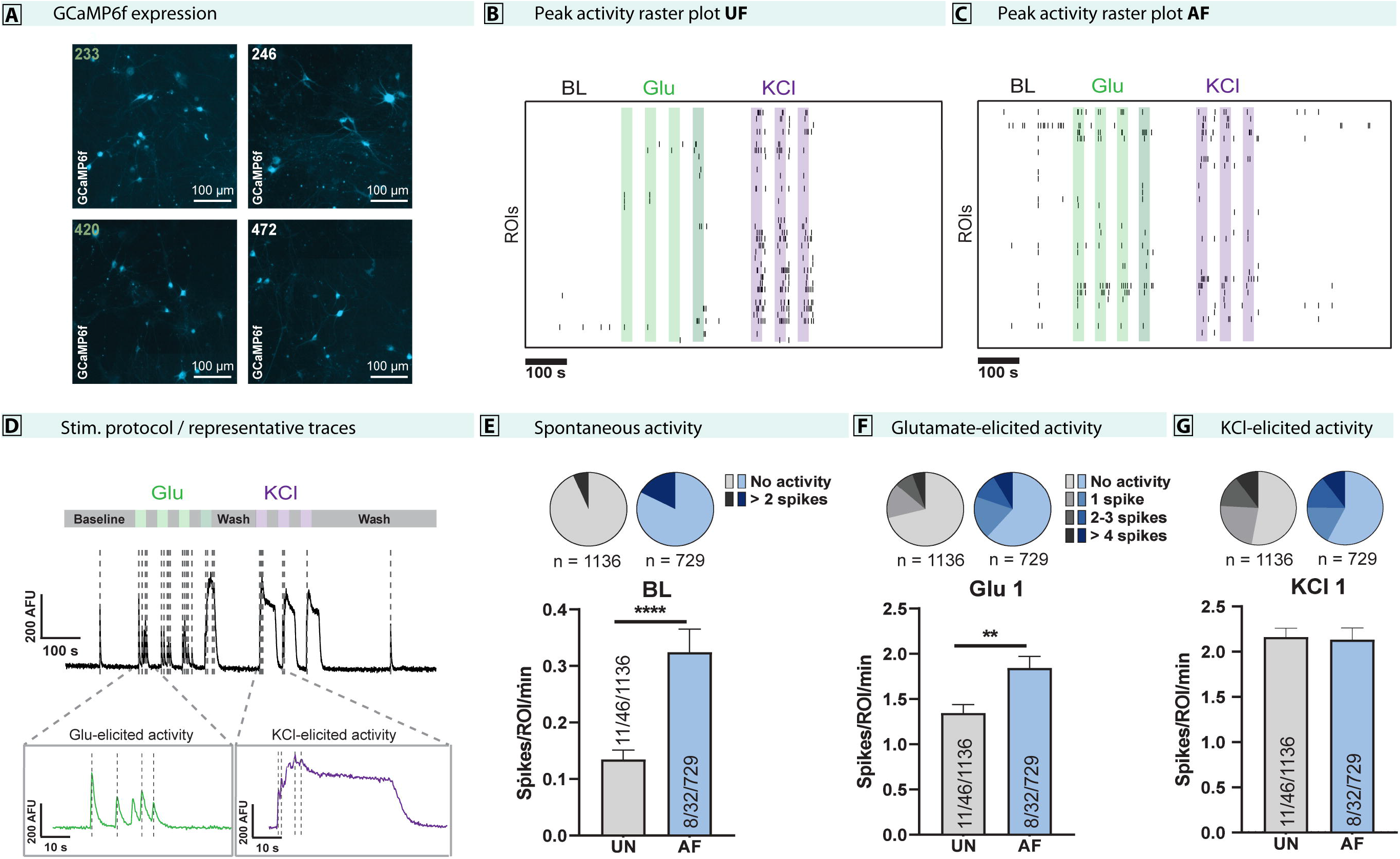
Evaluation of neuronal excitability in human induced neuron populations. **A**. Representative epifluorescence images of GcaMP6f expression in iNs from affected (233, 246) and unaffected individuals (420, 472). **B, C**. Raster plots of the calcium spiking pattern of representative experiments from unaffected neurons (**B**, individual 472) and affected neurons (**C**, individual 233), under the stimulation protocol used. Green and purple bars show the application of 30 s pulses of Glu 10-50 μM and KCl 18 mM, respectively, on the neuron populations during the stimulation protocol. Arrows show the epochs of the calcium imaging recordings selected for calcium spike quantification: baseline, glutamate pulse 1, and KCl pulse 1. **D**. Stimulation protocol (upper bar) and representative raw fluorescence trace (AF individual 233) from a complete recording. The zoom-ins on the first glutamate and KCl pulses depict glutamate and KCl elicited activity, respectively. The vertical dotted lines correspond to the calcium spikes detected after analysis. **E, F, G**. Number of spikes per ROI per minute during baseline (**E**, spontaneous activity), glutamate pulse 1 (**F**, glutamate-elicited activity) and KCl pulse 1 (**G**, KCl-elicited activity). The pie charts represent the proportion of neurons of UN (grey) and AF (blue) individuals that exhibited a certain number of calcium spikes during each three epochs of the recording. The bar plots show the number of spikes per ROI per minute fired by unaffected individuals (grey) and affected individuals (blue) during baseline, glutamate pulse 1 and KCl pulse 1. The sample size is depicted in the bar of each group as number of iN batches/number of experiments/number of neurons. Differences between affected and unaffected groups were evaluated by two-tailed unpaired Student’s t-test (Welch’s correction for non-equal SD, **p <0.01, ****p < 0.0001).

### Ethanol exposure eliminates differences in properties affected by *KCNJ6* haplotype

Analysis of *KCNJ6* mRNA expression in selected neurons (Fig. 1C) not only detected decreased basal levels in the AF group, but also demonstrated increased expression following ethanol treatment. This predicts that increasing GIRK2 expression in the AF group following ethanol treatment would eliminate differences from the UN group in morphology, expression, and physiology.

To control ethanol dosage in culture, we utilized an intermittent ethanol exposure (IEE) paradigm as in previous studies^25^. Ethanol was replenished daily to account for loss by evaporation, producing mean concentrations of 15.4 ± 1.16 mM (Supplemental Fig. 7). The half-life of ethanol is 14.5 h (with a 95% CI of 13.5 to 15.6 h), so over 24 h the concentration would drop to ^~^5-8 mM before replenishment. We targeted a peak concentration of 20 mM ethanol, which is similar to a blood alcohol concentration (BAC) of 0.08% (17 mM), the legal limit for intoxication. We also selected this concentration based on concentration-dependent interactions of GIRK2 with ethanol^17^. After 7 days of IEE, we evaluated neuronal morphological and functional properties. Focusing on parameters that were different by genotype without IEE (Fig. 3&4), we found that ethanol eliminated the differences in total neurite area (p = 0.98, Fig. 5A.d), GIRK2 puncta counts (p = 0.46, Fig. 5B.a), excitation following current injection (p = 0.32, Fig. 5E.a), step- (p = 0.48, Fig. 5E.b), and ramp- (p = 0.95, Fig. 5E.d) induced APs. Membrane capacitance was slightly but significantly reduced in the IEE AF group (p = 9.6 × 10^−9^, Fig. 5D.a). Other measurements were unchanged, again indicating that the cultured neurons exhibited similar differentiation properties. Results are consistent with the interpretation that neurite and excitability differences correlate with reduced GIRK2 expression, and that ethanol increases expression of GIRK2, reversing these effects.

**Figure 5.**
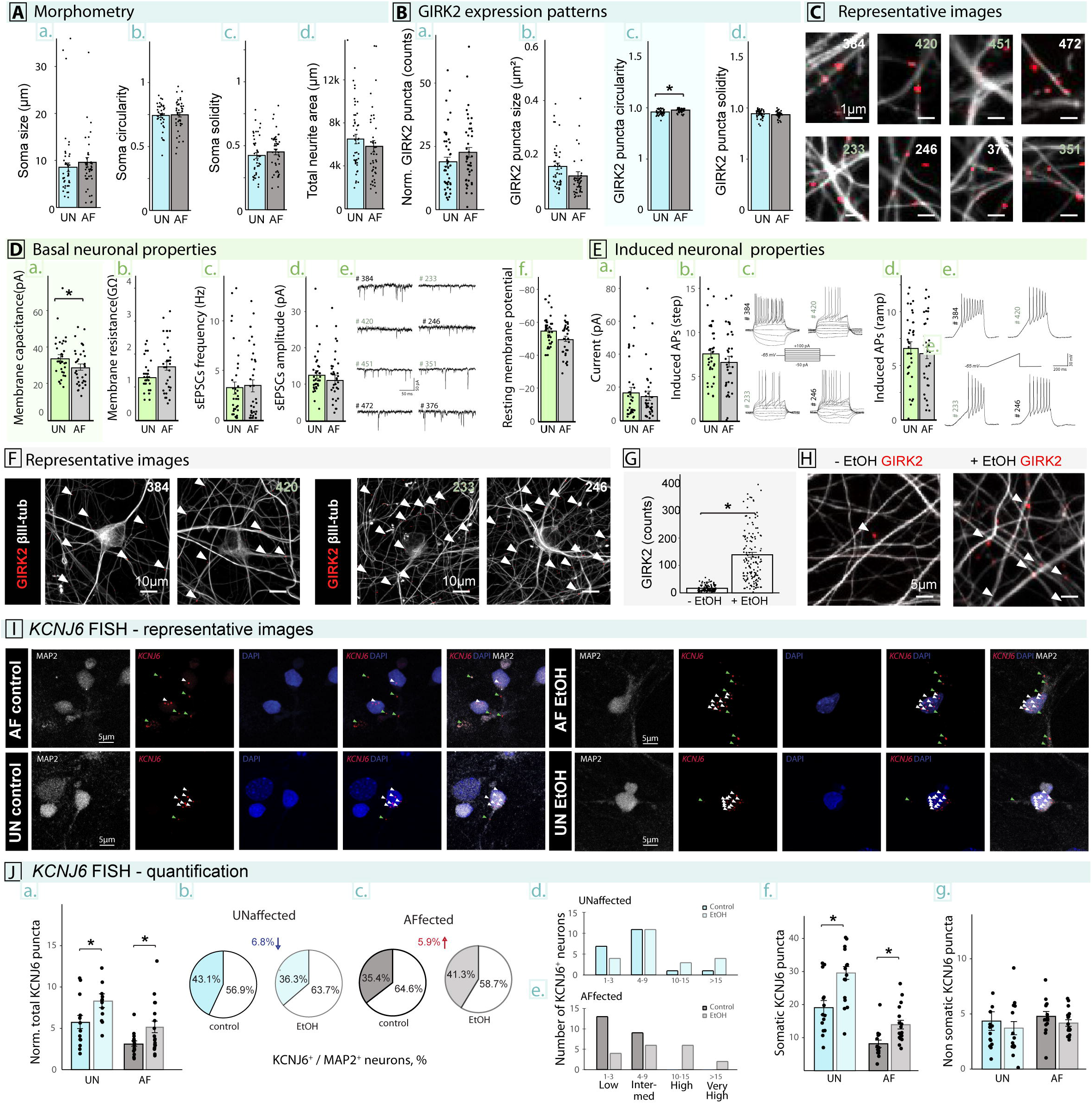
Ethanol treatment reduced *KCNJ6* haplotype differences in iN excitability and GIRK2 expression. **A**. Morphological analysis of IEE iNs generated from affected and unaffected individuals, showing no difference in: (a) neuronal soma size (p = 0.32); (b) soma circularity (p = 0.54); (c) soma solidity (p = 0.92); and (d) total neurite area (p = 0.98). **B**. There was no difference in GIRK2 expression in the IEE AF group iNs compared with UN by (a) puncta counts (p = 0.46), (b) puncta size (p = 0.36), (c) puncta circularity (p = 0.052), or (d) solidity (p = 0.47). **C**. Representative images of individual GIRK2 puncta (red) localized on βIII-Tubulin positive processes (gray) for each cell line. **D**. Electrophysiological analysis of passive neuronal properties in IEE iNs, showing (a) a small decrease in AF membrane capacitance (p = 9.6 × 10^−9^), but no difference in (b) membrane resistance (p = 0.63), (c) spontaneous EPSCs frequency (p = 0.32), or (d) spontaneous EPSCs amplitude (p = 0.19). (e) Representative sEPSCs traces for each cell line. (f) The AF group exhibited no change in resting membrane potential after IEE (p = 0.79). **E**. Electrophysiological analysis of active neuronal properties found no difference in IEE iNs for (a) current required to shift resting membrane potential to −65mV (p = 0.32), (b) maximum number of action potentials (APs) induced with the “step” protocol (p = 0.48), with (c) representative traces of APs induced with the “step” protocol, (d) number of action potentials (APs) induced with the “ramp” protocol (p = 0.95), with (e) representative traces of APs induced with the “ramp” protocol. **F**. Representative images from individual lines of iNs, exposed to 7d of IEE, marked with arrows pointing to individual GIRK2 puncta (red) localized on βIII-Tubulin positive processes (gray). **G**. Summarized results from all lines showing differences in GIRK2 expression levels before and after 7 days of 20 mM IEE with ethanol (EtOH; p = 1.0 × 10^−20^). **H**. Representative images of individual GIRK2 puncta (red) localized on βIII-Tubulin positive processes (gray) prior and following 7 days 20 mM IEE with ethanol. Sample images are from line 246. **I**. Representative images of FISH detection of *KCNJ6* mRNA for each cell line. **J**. Quantification of FISH. (a) The number of *KCNJ6* puncta normalized to the number of cells in an image shows decreased expression in control AF compared with UN (p = 8.2 × 10^−3^), increased expression following IEE (p = 8.2 × 10^−3^; using Tukey’s pairwise comparisons). (b) The percentage of *KCNJ6*-expressing MAP2^+^ cells substantially increase in the (c) AF group but not in the (b) UN group. Numbers of *KCNJ6* puncta were analyzed by expression levels per cell, as recommended by the FISH manufacturer in (d) UN or (e) AF cells. (f) *KCNJ6* puncta within the neuronal soma show increases following IEE in both UN and AF groups (p = 0.006 for genotype, Tukey’s post-hoc for UN, p = 0.01, for AF, p = 0.01). (g) A similar analysis of non-somatic puncta, presumably within neurites, showed no differences following IEE between genotypes (p=0.23).

To confirm that both *KCNJ6* haplotype and ethanol exposure affect GIRK2 expression, we evaluated GIRK2 immunocytochemistry and *KCNJ6* mRNA in neurons using fluorescent in situ hybridization (FISH). GIRK2 increased following IEE (p=1.0 × 10^−20^, Fig. 5G-H). By probing *KCNJ6* fluorescent puncta colocalized with MAP2 immunocytochemistry, we confirmed that neurons from AF individuals had lower levels of *KCNJ6* mRNA compared to UN neurons (Fig. 5J.a, p = 8.2 × 10^−3^). We also found that the proportion of MAP2^+^ neurons expressing *KCNJ6* was reduced in AF compared with UN (50.1% vs 64.7%, Fig. 5J.b), and the AF neurons had a greater proportion of cells with low levels of expression (Fig. 5J.c) with no detectable High or Very High (>9 puncta/cell) expression. Differences in expression were found by haplotype in somatic regions of neurons (Fig. 5J.f, p = 5.5 × 10^−3^) but not in non-somatic processes (Fig. 5J.g, p = 0.23). Ethanol increased *KCNJ6* mRNA levels in both groups (Fig. 5J.a-e), eliminating the difference between genotypes. Interestingly, we also observed that the proportions of AF neurons expressing *KCNJ6* was increased (8.3%, Fig. 5J.b) by ethanol and was paralleled by a reduction in number of low-expressing cells and increased appearance of high-expressing neurons (Fig. 5J.c). Results indicate that variant *KCNJ6* haplotype leads to reduced *KCNJ6* expression, which is consistent with RNA sequencing and immunocytochemical results, and that ethanol increases *KCNJ6* expression, eliminating differences between haplotype groups.

### GIRK2 overexpression mimics IEE in human neurons with *KCNJ6* variants

To test if ethanol-induced GIRK2 expression could underlie the elimination of differences in neuronal properties associated with the *KCNJ6* haplotype, we compared ethanol treatment with virus transduced *KCNJ6* overexpression. Since ethanol has been shown to potentiate GIRK2-containing channels, we used a single treatment with 20 mM ethanol followed by fixation 24 h later. As expected, neither ethanol nor GIRK2 overexpression exhibited differences in passive neuronal properties. However, while current-induced activity increased in control cultures (p = 6.8 × 10^−6^, Fig. 6C), no significant difference was found in cultures exposed to 24 h ethanol (p = 0.74) and GIRK2 overexpression (p = 0.23). Similarly, while control cultures had increased ramp-induced APs (p = 5.9 × 10^−4^; Fig. 6A) neither GIRK overexpression (p = 0.14) or ethanol (p = 0.10) were significantly different. As a control, GIRK2 puncta count increased in a sample cell line after overexpression (line 376, p = 0.04, Fig. 6D) without an increase in circularity or solidity (not shown). These results indicate that the increased excitability in *KCNJ6* minor allele haplotype cells is due to reduced expression of GIRK2 protein and this effect is at least partially ameliorated or reversed by exposure to doses of ethanol found in human brain after moderate to heavy daily drinking.

**Figure 6.**
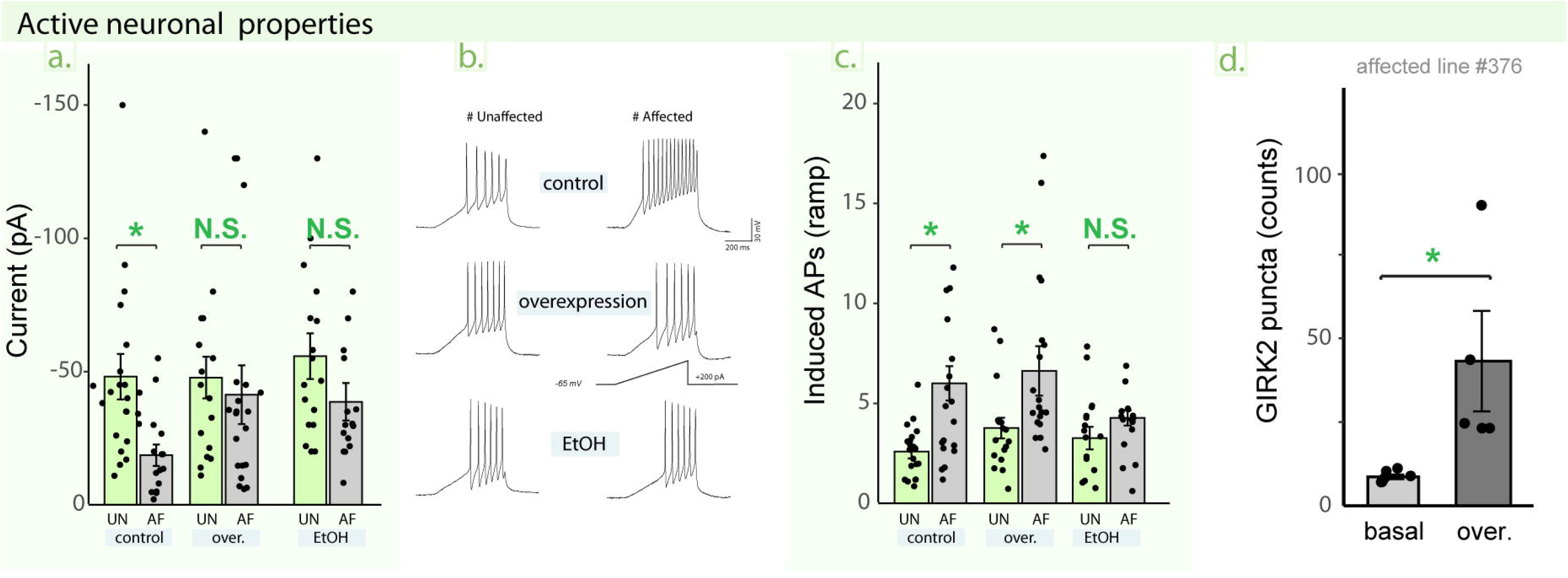
GIRK2 overexpression mimics ethanol response. **A**. Current required to shift resting membrane potential to −65mV, in untreated (control, p = 5.9 × 10^−4^), lentiviral *KCNJ6* overexpression (over., p = 0.23), or 1 d 20 mM IEE (EtOH, p = 0.75) cultures. **B**. Representative traces of APs induced with the “ramp” protocol. **C**. Quantification of maximum number of action potentials (APs) induced with “ramp” protocol, control (p = 6.8 × 10^−6^), overexpression (p = 0.014), or 1 d 20 mM IEE (p = 0.10). **D**. quantification of GIRK2 puncta after overexpression (line 376, one-tailed Student’s t-test p = 0.04).

## Discussion

This study focused on the effect of an AUD- and endophenotype-linked *KCNJ6* haplotype on neuronal function, with a goal of identifying mechanisms that could eventually drive therapeutic strategies. To retain the appropriate genetic background for individuals with AUD, we selected subjects from the NIAAA/COGA Sharing Repository, using both *KCNJ6* haplotype and alcohol dependence as criteria (Table 1). By preparing iPSC from these subjects and inducing them to model excitatory neurons, we identified differences in *KCNJ6* mRNA levels and GIRK2 immunoreactivity in individuals with variant *KCNJ6* haplotypes. Gene expression differences predicted effects on neuronal signaling including synaptic function. The *KCNJ6* haplotype was also associated with differences in neuron projection area and membrane excitability, but not in other measures of neuronal differentiation. Surprisingly, we found that ethanol exposure led to increased *KCNJ6* mRNA and GIRK2 expression, paralleled by reduced or eliminated morphological and physiological differences. Reversal of *KCNJ6* haplotype effects by ethanol could also be mimicked by overexpressing GIRK2. These results demonstrate that genetic variants enhancing risk of AUD, even if they do not alter protein sequence, can trigger neuronal mechanisms at the cellular level mirroring or predicting endophenotypes, here ERO, linked with AUD behavior.

### Non-coding *KCNJ6* polymorphisms in AUD

Most GWAS SNPs map to non-coding regions of the genome and 3.7% are found in the UTRs^47,48^. The 3Ͱ UTRs of transcripts expressed in brain tend to be longer than those expressed in other tissues and can be involved in post-transcriptional regulation of transcript abundance by affecting RNA stability, translation and/or localization^49,50^. Annotation of *KCNJ6* identifies two transcripts, one with a short 3’ UTR (1,926 bp; ENST00000645093) and one with a substantially longer 3’ UTR (18,112 bp; ENST00000609713). RNAseq coverage analysis in human iN detects only the longer isoform, although it is possible that a relatively small portion could be the shorter isoform, but the uniformly distributed coverage (Fig. 1B) does not indicate this. This longer transcript is orthologous to a 16-kb transcript found in rat brain^51^, which includes multiple AU-rich elements, which may affect mRNA stability and could be affected by *KCNJ6* allelic variation. We conclude that the vast majority of *KCNJ6* mRNA is consistent with having the extended 3’ UTR, with the potential for variant SNPs to affect mRNA stability or translation efficiency.

In addition to the three synonymous and intronic SNPs used to select subjects (Table 1), examination of variants in the long 3’UTR by RNAseq alignment predicts linkage of 19 additional SNPs (Fig. 1B; Supplemental Table 1). None of the 3’ UTR SNPs could be mapped uniquely to known regulatory sequences such as predicted microRNA target sites. That is, some SNPs are predicted to destroy or add predicted targeting sites, but all were found in multiple locations within the 3’ UTR, not only where they might be altered by a SNP. It is possible, however, that altered microRNA targeting of one or more sites among many in the 3’UTR could play a role in regulating the stability or translation of the variant *KCNJ6* mRNA, but no clear candidates could be identified for testing.

However, results demonstrate that the non-coding haplotype alters expression of GIRK2 protein and *KCNJ6* mRNA. iN from subjects with the homozygous variant *KCNJ6* haplotype exhibited lower expression of GIRK2 by immunocytochemistry (Fig. 3D), but while the difference in neuronal mRNA levels as detected by scRNAseq analysis matched this trend, it did not reach significance (p = 0.0508; Fig. 1C). FISH analysis, however, confirmed that *KCNJ6* mRNA was reduced in the AF group (Fig. 5J). *KCNJ6* mRNA was also detected in fewer MAP2^+^ cells and at lower levels per cell (Fig. 5J.b-c). The results of counting GIRK2 puncta and evaluating *KCNJ6* mRNA by scRNAseq and FISH support a diminished expression of GIRK2 in neurons from the affected group. Secondary effects of *KCNJ6* mRNA and GIRK2 protein regulation include many predictors of altered neuronal connectivity and signaling. Genes that were reduced in the AF group (GO:BP Down in Fig. 1E; Supplemental Fig. 3B) point to changes in synaptic transmission and projection morphology. Since we focused on *KCNJ6* haplotype differences in cells without ethanol, we did not expect to match findings in, for example, human AUD brains^52^, which found prominent effects in astrocytes and microglia.

Surprisingly, ethanol exposure increased expression of both GIRK2 immunoreactivity and *KCNJ6* mRNA. Following 7 days of IEE, GIRK2 puncta in the AF group increased to levels significantly higher than the UN group (Fig. 5B). Similarly, *KCNJ6* mRNA levels were shown to be increased by scRNAseq results (Fig. 1C) and FISH analysis (Fig. 5J). While previous studies identified a binding site for ethanol in a defined pocket of GIRK2^53^, suggesting that it could affect protein stabilization, this could not explain the increased mRNA level. There is no indication that ethanol or *KCNJ6* haplotype altered translocation of *KCNJ6* mRNA to neuronal processes (Fig. 5J.d-e). Ethanol has been found to affect microRNA expression patterns that regulate mRNA^49,54–59^. It is intriguing to speculate that ethanol might increase expression of GIRK2 through this type of mechanism, with polymorphisms in the 3’UTR serving as the modulator.

### GIRK2 in the context of alcohol dependence

GIRK channels play an essential role in maintaining the excitability of neurons. For example, in Trisomy 21 individuals, whose cells contain an additional copy of *KCNJ6*, neuronal activity was shown to be reduced due to excessive channel activity^60–62^. On the other hand, loss of GIRK2 activity in knockout mice increases susceptibility to induced seizures^20,21^. We detected the presence of GIRK channel function in glutamatergic iNs using the ML297 agonist of GIRK1/2 heterotetramers, showing that activation led to a reduction in the frequency of action potentials induced by depolarization (Fig. 2D). However, only 6.7% of neurons responded to ML297, but overexpression of GIRK2 increased the proportion of responding neurons to 30% (Fig. 2D). We therefore chose to focus on evaluating neuronal excitability as an indirect response to changes in GIRK function.

By identifying the correlation between GIRK2 expression levels and altered GIRK2-mediated function, we predict that neuronal excitability will be affected by *KCNJ6* variants and alcohol exposure. While GIRK channels play relatively small role in maintaining neuronal homeostasis, modulating GIRK activity to alter cell excitability is predicted to play a critical role. Additionally, downstream effectors of the G protein-mediated signaling pathways presumably regulate neuronal function indirectly^14,15,17,19^. GIRK channels are implicated in several other disorders with abnormal neuronal excitability, including epilepsy, suggesting that they have therapeutic potential^19–21,60–63^.

### Potential limitations

Factors beyond *KCNJ6* haplotype may contribute to these interpretations. It is possible that the sex of individual neurons might contribute to the results. However, in nearly all parameters tested, there was no difference by sex. scRNAseq identified only 6 genes as different by sex and 4 of these are encoded by sex chromosomes. Basal morphology (Fig. 3B, D) was unaffected by sex, but membrane capacitance (Fig. 3F.a, p = 2.6 × 10^−4^), induced APs (step, Fig. 3G.b, p = 0.012), and ethanol-treated membrane capacitance (Fig. 5D.a, p = 3.1 × 10^−8^) all reached significance, although it is difficult to interpret how sex interacts with neuronal function in this context. Finally, we chose to focus on homozygous haplotypes and extreme AUD diagnoses to enhance contrasts, however, additional studies might examine if heterozygotes would exhibit gene dosage effects or be rescued by the presence of the major alleles, similar to endophenotype studies with the original subjects^8^.

### From genes to behavior

In this study, we found that the variant *KCNJ6* haplotype affects excitability of neurons (Fig. 3 & 4E). In Ca^2+^ imaging experiments, concomitant with enhanced excitability in neurons from affected individuals following stimulation with glutamate pulses, we observed an overall increase in basal activity of the neuronal population (Fig. 4F). This is reminiscent of the original observation linking *KCNJ6* variants with an EEG endophenotype, where Kamarajan and colleagues observed that individuals with alcohol dependence and the *KCNJ6* haplotype had an ERO theta power that varied as a function of the *KCNJ6* haplotype in both loss and gain conditions^10^. There is an extensive human literature linking impulsivity to alcohol use problems^64–67^. Impulsivity is elevated in offspring who are at high risk for substance use disorders and may be a reflection of a genetic vulnerability for substance use problems^68^. Excitability in individual neurons and particularly in a culture dish of neurons is several levels of complexity removed from EEG patterns in brain. However, our study points to a mechanistic underpinning of how heritable risk traits are likely to play a role in development of unique physiological response to alcohol in the brain.

## Acknowledgements

The Collaborative Study on the Genetics of Alcoholism (COGA), Principal Investigators B. Porjesz, V. Hesselbrock, T. Foroud; Scientific Director, A. Agrawal; Translational Director, D. Dick, includes eleven different centers: University of Connecticut (V. Hesselbrock); Indiana University (H.J. Edenberg, T. Foroud, Y. Liu, M. Plawecki); University of Iowa Carver College of Medicine (S. Kuperman, J. Kramer); SUNY Downstate Health Sciences University (B. Porjesz, J. Meyers, C. Kamarajan, A. Pandey); Washington University in St. Louis (L. Bierut, J. Rice, K. Bucholz, A. Agrawal); University of California at San Diego (M. Schuckit); Rutgers University (J. Tischfield, R. Hart, J. Salvatore); The Children’s Hospital of Philadelphia, University of Pennsylvania (L. Almasy); Virginia Commonwealth University (D. Dick); Icahn School of Medicine at Mount Sinai (A. Goate, P. Slesinger); and Howard University (D. Scott). Other COGA collaborators include: L. Bauer (University of Connecticut); J. Nurnberger Jr., L. Wetherill, X., Xuei, D. Lai, S. O’Connor, (Indiana University); G. Chan (University of Iowa; University of Connecticut); D.B. Chorlian, J. Zhang, P. Barr, S. Kinreich, G. Pandey (SUNY Downstate); N. Mullins (Icahn School of Medicine at Mount Sinai); A. Anokhin, S. Hartz, E. Johnson, V. McCutcheon, S. Saccone (Washington University); J. Moore, Z. Pang, S. Kuo (Rutgers University); A. Merikangas (The Children’s Hospital of Philadelphia and University of Pennsylvania); F. Aliev (Virginia Commonwealth University); H. Chin and A. Parsian are the NIAAA Staff Collaborators. We continue to be inspired by our memories of Henri Begleiter and Theodore Reich, founding PI and Co-PI of COGA, and also owe a debt of gratitude to other past organizers of COGA, including Ting-Kai Li, P. Michael Conneally, Raymond Crowe, and Wendy Reich, for their critical contributions. This national collaborative study is supported by NIH Grant U10AA008401 from the National Institute on Alcohol Abuse and Alcoholism (NIAAA) and the National Institute on Drug abuse (NIDA).

## Conflict of Interest

The authors declare no conflict of interest.

## Supplementary Materials

**Supplemental Tables 1-6: https://doi.org/10.6084/m9.figshare.19798882.v2**

**Supplemental Table 1. SNPs identified in RNAseq samples**. Variants were identified in the region surrounding KCNJ6 (chr21: 37499112-38245792) using samtools mpileup to bcftools call, followed by filtering to remove low quality. Results were loaded into the Ensembl VEP tool. A dot (“.”) indicates missing data and/or low quality. The reference allele and alternate alleles were obtained from the UCSC Genome Table Browser.

**Supplemental Table 2: CellRanger parameters for scRNAseq libraries**. Output from CellRanger shows the alignment statistics and numbers of reads confidently mapped.

**Supplemental Table 3. Differentially expressed genes (DEG) comparing AF to UN without exposure to ethanol**. The table contains all output from DESeq2 results. Excel filters are set to show only genes significantly different (padj <= 0.05) and at least 1.5-fold different (abs(log2FoldChange) > 0.585). Release the filters to see the full list.

**Supplemental Table 4. Enriched Gene Ontology-Biological Process (GO-BP) terms from genes increased in AF relative to UN**.

**Supplemental Table 5. Enriched Gene Ontology-Biological Process (GO-BP) terms from genes decreased in AF relative to UN**.

**Supplemental Table 6. Numbers of replicates**. For each parameter tested with statistics, the numbers of cells and/or microscope fields is indicated and labeled by Figure numbers/letters.

**Supplemental Methods and Figures: https://doi.org/10.6084/m9.figshare.19798873.v5**

## Notes

https://doi.org/10.6084/m9.figshare.19798873.v5

https://doi.org/10.6084/m9.figshare.19798882.v2

### Competing Interest Statement

The authors have declared no competing interest.

### Summary of Updates

Updated version with new supplemental methods cited.

## References

1. Verhulst B, Neale MC, Kendler KS. The heritability of alcohol use disorders: a meta-analysis of twin and adoption studies. Psychol Med 2015; 45(5):1061–1072.

2. Pollard MS, Tucker JS, Green HD, Jr. Changes in Adult Alcohol Use and Consequences During the COVID-19 Pandemic in the US. JAMA Network Open 2020; 3(9):e2022942–e2022942.

3. Bouza C, Angeles M, Muñoz A, Amate JM. Efficacy and safety of naltrexone and acamprosate in the treatment of alcohol dependence: a systematic review. Addiction 2004; 99(7):811–828.

4. Heilig M, Egli M. Pharmacological treatment of alcohol dependence: Target symptoms and target mechanisms. Pharmacology & Therapeutics 2006; 111(3):855–876.

5. Skinner MD, Lahmek P, Pham H, Aubin HJ. Disulfiram efficacy in the treatment of alcohol dependence: a meta-analysis. PLoS One 2014; 9(2):e87366.

6. Salvatore JE, Gottesman, II, Dick DM. Endophenotypes for Alcohol Use Disorder: An Update on the Field. Curr Addict Rep 2015; 2(1):76–90.

7. Chorlian DB, Rangaswamy M, Manz N, Meyers JL, Kang SJ, Kamarajan C et al. Genetic correlates of the development of theta event related oscillations in adolescents and young adults. Int J Psychophysiol 2017; 115:24–39.

8. Kang SJ, Rangaswamy M, Manz N, Wang JC, Wetherill L, Hinrichs T et al. Family-based genome-wide association study of frontal θ oscillations identifies potassium channel gene KCNJ6. Genes Brain Behav 2012; 11(6):712–719.

9. Clarke TK, Laucht M, Ridinger M, Wodarz N, Rietschel M, Maier W et al. KCNJ6 is associated with adult alcohol dependence and involved in gene × early life stress interactions in adolescent alcohol drinking. Neuropsychopharmacology 2011; 36(6):1142–1148.

10. Kamarajan C, Pandey AK, Chorlian DB, Manz N, Stimus AT, Edenberg HJ et al. A KCNJ6 gene polymorphism modulates theta oscillations during reward processing. Int J Psychophysiol 2017; 115:13–23.

11. Andrew C, Fein G. Event-related oscillations versus event-related potentials in a P300 task as biomarkers for alcoholism. Alcohol Clin Exp Res 2010; 34(4):669–680.

12. Kamarajan C, Rangaswamy M, Tang Y, Chorlian DB, Pandey AK, Roopesh BN et al. Dysfunctional reward processing in male alcoholics: an ERP study during a gambling task. J Psychiatr Res 2010; 44(9):576–590.

13. Glaaser IW, Slesinger PA. Structural Insights into GIRK Channel Function. International review of neurobiology 2015; 123:117–160.

14. Luscher C, Slesinger PA. Emerging roles for G protein-gated inwardly rectifying potassium (GIRK) channels in health and disease. Nat Rev Neurosci 2010; 11(5):301–315.

15. Zhao Y, Gameiro-Ros I, Glaaser IW, Slesinger PA. Advances in Targeting GIRK Channels in Disease. Trends Pharmacol Sci 2021; 42(3):203–215.

16. Reuveny E, Slesinger PA, Inglese J, Morales JM, Iñiguez-Lluhi JA, Lefkowitz RJ et al. Activation of the cloned muscarinic potassium channel by G protein beta gamma subunits. Nature 1994; 370(6485):143–146.

17. Aryal P, Dvir H, Choe S, Slesinger PA. A discrete alcohol pocket involved in GIRK channel activation. Nat Neurosci 2009; 12(8):988–995.

18. Bodhinathan K, Slesinger PA. Alcohol modulation of G-protein-gated inwardly rectifying potassium channels: from binding to therapeutics. Front Physiol 2014; 5:76.

19. Blednov YA, Stoffel M, Chang SR, Harris RA. Potassium channels as targets for ethanol: studies of G-protein-coupled inwardly rectifying potassium channel 2 (GIRK2) null mutant mice. J Pharmacol Exp Ther 2001; 298(2):521–530.

20. Blednov YA, Stoffel M, Chang SR, Harris RA. GIRK2 deficient mice. Evidence for hyperactivity and reduced anxiety. Physiol Behav 2001; 74(1-2):109–117.

21. Hill KG, Alva H, Blednov YA, Cunningham CL. Reduced ethanol-induced conditioned taste aversion and conditioned place preference in GIRK2 null mutant mice. Psychopharmacology (Berl) 2003; 169(1):108–114.

22. Lieberman R, Levine ES, Kranzler HR, Abreu C, Covault J. Pilot study of iPS-derived neural cells to examine biologic effects of alcohol on human neurons in vitro. Alcohol Clin Exp Res 2012; 36(10):1678–1687.

23. Lieberman R, Kranzler HR, Levine ES, Covault J. Examining the effects of alcohol on GABA(A) receptor mRNA expression and function in neural cultures generated from control and alcohol dependent donor induced pluripotent stem cells. Alcohol 2018; 66:45–53.

24. Halikere A, Popova D, Scarnati MS, Hamod A, Swerdel MR, Moore JC et al. Addiction associated N40D mu-opioid receptor variant modulates synaptic function in human neurons. Mol Psychiatry 2020; 25(7):1406–1419.

25. Scarnati MS, Boreland AJ, Joel M, Hart RP, Pang ZP. Differential sensitivity of human neurons carrying mu opioid receptor (MOR) N40D variants in response to ethanol. Alcohol 2020; 87:97–109.

26. Patzke C, Dai J, Brockmann MM, Sun Z, Fenske P, Rosenmund C et al. Cannabinoid receptor activation acutely increases synaptic vesicle numbers by activating synapsins in human synapses. Mol Psychiatry 2021.

27. Vierbuchen T, Ostermeier A, Pang ZP, Kokubu Y, Sudhof TC, Wernig M. Direct conversion of fibroblasts to functional neurons by defined factors. Nature 2010; 463(7284):1035–1041.

28. Pang ZP, Yang N, Vierbuchen T, Ostermeier A, Fuentes DR, Yang TQ et al. Induction of human neuronal cells by defined transcription factors. Nature 2011; 476(7359):220–223.

29. Zhang Y, Pak C, Han Y, Ahlenius H, Zhang Z, Chanda S et al. Rapid single-step induction of functional neurons from human pluripotent stem cells. Neuron 2013; 78(5):785–798.

30. Lin HC, He Z, Ebert S, Schörnig M, Santel M, Nikolova MT et al. NGN2 induces diverse neuron types from human pluripotency. Stem Cell Reports 2021; 16(9):2118–2127.

31. Bardy C, van den Hurk M, Eames T, Marchand C, Hernandez RV, Kellogg M et al. Neuronal medium that supports basic synaptic functions and activity of human neurons in vitro. Proc Natl Acad Sci U S A 2015; 112(20):E2725–2734.

32. Schneider CA, Rasband WS, Eliceiri KW. NIH Image to ImageJ: 25 years of image analysis. Nature methods 2012; 9(7):671–675.

33. Fantuzzo JA, Robles DA, Mirabella VR, Hart RP, Pang ZP, Zahn JD. Development of a high-throughput arrayed neural circuitry platform using human induced neurons for drug screening applications. Lab Chip 2020; 20(6):1140–1152.

34. Bates D, Mächler M, Bolker B, Walker S. Fitting Linear Mixed-Effects Models Using lme4. Journal of Statistical Software 2015; 67(1):1 – 48.

35. Højsgaard S, Halekoh U, Yan J. The R Package geepack for Generalized Estimating Equations. Journal of Statistical Software 2005; 15(2):1 – 11.

36. Love MI, Huber W, Anders S. Moderated estimation of fold change and dispersion for RNA-seq data with DESeq2. Genome Biol 2014; 15(12):550.

37. Mitchell JM, Nemesh J, Ghosh S, Handsaker RE, Mello CJ, Meyer D et al. Mapping genetic effects on cellular phenotypes with “cell villages”. bioRxiv 2020:2020.2006.2029.174383.

38. Lesage F, Guillemare E, Fink M, Duprat F, Heurteaux C, Fosset M et al. Molecular properties of neuronal G-protein-activated inwardly rectifying K+ channels. J Biol Chem 1995; 270(48):28660–28667.

39. Ang CE, Wernig M. Induced neuronal reprogramming. J Comp Neurol 2014; 522(12):2877–2886.

40. Shelby H, Shelby T, Wernig M. Somatic Lineage Reprogramming. Cold Spring Harb Perspect Biol 2021.

41. Kang HM, Subramaniam M, Targ S, Nguyen M, Maliskova L, McCarthy E et al. Multiplexed droplet single-cell RNA-sequencing using natural genetic variation. Nat Biotechnol 2018; 36(1):89–94.

42. Marron Fernandez de Velasco E, Zhang L, B NV, Tipps M, Farris S, Xia Z et al. GIRK2 splice variants and neuronal G protein-gated K(+) channels: implications for channel function and behavior. Sci Rep 2017; 7(1):1639.

43. Wydeven N, Marron Fernandez de Velasco E, Du Y, Benneyworth MA, Hearing MC, Fischer RA et al. Mechanisms underlying the activation of G-protein-gated inwardly rectifying K+ (GIRK) channels by the novel anxiolytic drug, ML297. Proc Natl Acad Sci U S A 2014; 111(29):10755–10760.

44. Days E, Kaufmann K, Romaine I, Niswender C, Lewis M, Utley T et al. Discovery and Characterization of a Selective Activator of the G-Protein Activated Inward-Rectifying Potassium (GIRK) Channel. Probe Reports from the NIH Molecular Libraries Program. National Center for Biotechnology Information (US): Bethesda (MD), 2010.

45. Kang S, Chen X, Gong S, Yu P, Yau S, Su Z et al. Characteristic analyses of a neural differentiation model from iPSC-derived neuron according to morphology, physiology, and global gene expression pattern. Scientific reports 2017; 7(1):12233.

46. Chen TW, Wardill TJ, Sun Y, Pulver SR, Renninger SL, Baohan A et al. Ultrasensitive fluorescent proteins for imaging neuronal activity. Nature 2013; 499(7458):295–300.

47. Steri M, Idda ML, Whalen MB, Orrù V. Genetic variants in mRNA untranslated regions. Wiley Interdiscip Rev RNA 2018; 9(4):e1474.

48. Wang D, Liu S, Warrell J, Won H, Shi X, Navarro FCP et al. Comprehensive functional genomic resource and integrative model for the human brain. Science 2018; 362(6420).

49. Nunez YO, Mayfield RD. Understanding Alcoholism Through microRNA Signatures in Brains of Human Alcoholics. Front Genet 2012; 3:43.

50. Wehrspaun C, Ponting C, Marques A. Brain-expressed 3’UTR extensions strengthen miRNA cross-talk between ion channel/transporter encoding mRNAs. Frontiers in Genetics 2014; 5(41).

51. Suda S, Nibuya M, Suda H, Takamatsu K, Miyazaki T, Nomura S et al. Potassium channel mRNAs with AU-rich elements and brain-specific expression. Biochem Biophys Res Commun 2002; 291(5):1265–1271.

52. Brenner E, Tiwari GR, Kapoor M, Liu Y, Brock A, Mayfield RD. Single cell transcriptome profiling of the human alcohol-dependent brain. Hum Mol Genet 2020; 29(7):1144–1153.

53. Bodhinathan K, Slesinger PA. Molecular mechanism underlying ethanol activation of G-protein-gated inwardly rectifying potassium channels. Proc Natl Acad Sci U S A 2013; 110(45):18309–18314.

54. Sathyan P, Golden HB, Miranda RC. Competing interactions between micro-RNAs determine neural progenitor survival and proliferation after ethanol exposure: evidence from an ex vivo model of the fetal cerebral cortical neuroepithelium. J Neurosci 2007; 27(32):8546–8557.

55. Pietrzykowski AZ, Friesen RM, Martin GE, Puig SI, Nowak CL, Wynne PM et al. Posttranscriptional regulation of BK channel splice variant stability by miR-9 underlies neuroadaptation to alcohol. Neuron 2008; 59(2):274–287.

56. Lewohl JM, Nunez YO, Dodd PR, Tiwari GR, Harris RA, Mayfield RD. Up-regulation of microRNAs in brain of human alcoholics. Alcohol Clin Exp Res 2011; 35(11):1928–1937.

57. Osterndorff-Kahanek EA, Tiwari GR, Lopez MF, Becker HC, Harris RA, Mayfield RD. Long-term ethanol exposure: Temporal pattern of microRNA expression and associated mRNA gene networks in mouse brain. PLoS One 2018; 13(1):e0190841.

58. Lim Y, Beane-Ebel JE, Tanaka Y, Ning B, Husted CR, Henderson DC et al. Exploration of alcohol use disorder-associated brain miRNA-mRNA regulatory networks. Translational psychiatry 2021; 11(1):504.

59. Zhu S, Wu J, Hu J. Non-coding RNA in alcohol use disorder by affecting synaptic plasticity. Exp Brain Res 2022.

60. Best TK, Siarey RJ, Galdzicki Z. Ts65Dn, a mouse model of Down syndrome, exhibits increased GABAB-induced potassium current. J Neurophysiol 2007; 97(1):892–900.

61. Harashima C, Jacobowitz DM, Witta J, Borke RC, Best TK, Siarey RJ et al. Abnormal expression of the G-protein-activated inwardly rectifying potassium channel 2 (GIRK2) in hippocampus, frontal cortex, and substantia nigra of Ts65Dn mouse: a model of Down syndrome. J Comp Neurol 2006; 494(5):815–833.

62. Reeves RH, Irving NG, Moran TH, Wohn A, Kitt C, Sisodia SS et al. A mouse model for Down syndrome exhibits learning and behaviour deficits. Nat Genet 1995; 11(2):177–184.

63. Kleschevnikov AM, Yu J, Kim J, Lysenko LV, Zeng Z, Yu YE et al. Evidence that increased Kcnj6 gene dose is necessary for deficits in behavior and dentate gyrus synaptic plasticity in the Ts65Dn mouse model of Down syndrome. Neurobiol Dis 2017; 103:1–10.

64. Congdon E, Canli T. The endophenotype of impulsivity: reaching consilience through behavioral, genetic, and neuroimaging approaches. Behav Cogn Neurosci Rev 2005; 4(4):262–281.

65. Dick DM, Smith G, Olausson P, Mitchell SH, Leeman RF, O’Malley SS et al. Understanding the construct of impulsivity and its relationship to alcohol use disorders. Addict Biol 2010; 15(2):217–226.

66. Sher KJ, Trull TJ. Personality and disinhibitory psychopathology: alcoholism and antisocial personality disorder. J Abnorm Psychol 1994; 103(1):92–102.

67. Verdejo-García A, Lawrence AJ, Clark L. Impulsivity as a vulnerability marker for substance-use disorders: review of findings from high-risk research, problem gamblers and genetic association studies. Neurosci Biobehav Rev 2008; 32(4):777–810.

68. Polich J, Bloom FE. P300, alcoholism heritability, and stimulus modality. Alcohol 1999; 17(2):149–156.

69. Cáceres A, Banker GA, Binder L. Immunocytochemical localization of tubulin and microtubule-associated protein 2 during the development of hippocampal neurons in culture. J Neurosci 1986; 6(3):714–722.

70. Ma F, Xu J, Liu Y, Popova D, Youssef MM, Hart RP et al. The amyloid precursor protein modulates the position and length of the axon initial segment offering a new perspective on Alzheimer’s disease genetics. bioRxiv 2022:2022.2001.2023.477413.

