## Supplemental Methods and Figures for "Alcohol reverses the effects of *KCNJ6* (GIRK2) noncoding variants on excitability of human glutamatergic neurons"

### Supplementary Methods

#### Generation of human iPS cells and glutamatergic neurons (iN)

Cryopreserved lymphocytes were reprogrammed to iPSC using Sendai-viral-expressed transcription factors (Oct3/4, Sox2, Klf4 and c-Myc; Cytotune™-iPS Reprogramming Kit, ThermoFisher) by RUCDR Infinite Biologics® (now Sampled, Inc.). Pluripotent cells were selected by colony morphology, alkaline phosphatase positivity, and immunocytochemical expression of Oct4 and TRA-1-60. Pluripotency was confirmed by colony morphology and immunocytochemistry for Oct4 and Tra-1-60 markers (Supplemental Fig. 1A). Identity of each iPSC line was confirmed by matching a 96 SNP panel to results obtained from fresh blood upon banking the original sample. Cultures were tested for mycoplasma (all negative) prior to release by RUCDR. Identities of the SNPs listed in Table 1 were confirmed in all iPSC lines by PCR amplification from genomic DNA and Sanger sequencing. Genomic integrity was confirmed using e-Karyotyping[1] using RNAseq data, which detected no anomalies (Supplemental Fig. 1B). Frozen aliquots were obtained for each subject, thawed, expanded, and maintained for neurogenesis. All iPS lines were cultured up to 50 passages for these experiments.

#### Induced neuron culture

Briefly, iPS cells were plated as dissociated cells on Matrigel® Matrix (Corning Life Sciences)-coated dishes in iPS-Brew-XF (StemMACS) medium supplemented with 2 µM Y-27632 (Peprotech, Inc.) and 2 ng/µl doxycycline, and infected with lentiviruses expressing Ngn2 (AddGene #52047) and rtTA (FUW-M2rtTA; AddGene # 20342) for 10−12h. The next day, the culture medium was replaced with Neurobasal medium/B27/GlutaMAX™ (Life Technologies) supplemented with 2 µg/mL of doxycycline (MP Biomedical), 2 µM Y-27632 and 2 ng/µl puromycin. Puromycin selection was continued for 2 days, and on day 5, the induced neurons were dissociated with Accutase (STEMCELL Technologies) and plated on Matrigel-covered glass coverslips with a monolayer of primary astrocytes (passage 2-3) isolated from postnatal day 0-1 mouse pups, as described previously[2]. Following plating, 50% of Neurobasal medium was changed to Neurobasal Plus culture medium with ascorbic acid, GlutaMAX, B27 Plus and Culture One supplements (Life Technologies). Half of the Neurobasal Plus medium with all supplements was replaced every 2−3 days.

#### Primary mouse cortical neuron cultures

P0-P1 mouse brains were obtained following decapitation on ice, meninges were removed, cortex was dissected and cells were dissociated by incubation in trypsin (Gibco) for 5-7 min at 37°C. The cells were then washed with Hank's balanced salt solution (HBS) and triturated by gentle pipetting. The cell suspension was centrifuged for 5 min at 160 x g and plated on Matrigel-coated glass coverslips in Neurobasal medium/B27/L-Glutamine (Life Technologies). When the density of glia in the culture reached ∼40–50% (approximately 2 days after plating), 50% of the conditioned culture medium was replaced with fresh medium containing 4 mM Ara-C (Sigma; 2 mM final concentration). The cultures were maintained in medium containing 2 mM Ara-C until fixation at 13–18 days in vitro (DIV).

### Generation of CRISPR/Cas9 Knockout iPSC

As a negative control for GIRK2 antibody validation we generated a *KCNJ6* knockout by CRISPR/Cas9 deletion in WTC-11, a publicly-available, control iPS line obtained from the Gladstone Institute[3]. Two optimal gRNA targeting sequences were selected (using http://crispr.mit.edu) to generate a frameshift 76-bp deletion, introducing three termination codons within 40 bp following the deletion. A well-characterized iPSC line, WTB[4], was obtained from the Gladstone Institute Stem Cell Core Facility (San Francisco, CA). Approximately 3 x $10^6$ WTB iPSC (passage 58) were electroporated (amaxa Nucleofector) in suspension with a mixture of two plasmids constructed from pSpCas9(BB)-T2A-GFP [AddGene #48138[5]], with sequences targeting *KCNJ6*: G1, 5'-GATATCGGTCAGGTAGCGAT-3'; or G2, 5'-TCAGCCGAGATCGGACCAAA-3'. One day after transfection, cells were dissociated with Accutase and sorted to select $GFP^+$ cells into KSR medium (Life Technologies) with 2 µM Y-27632 and FGF (Peprotech, Inc.) and then plated at low density into two wells of a 6-well plate. Approximately 7-10 days later, individual colonies were hand-picked into 96-well plates. Three days after plating, cultures were split to two 96-well plates. One plate was harvested at ~50-100% confluence for screening. Cells were lysed in Quickextract (Lucigen), heated to 68°C for 6 minutes followed by 98°C for 2 minutes. The lysate was PCR amplified (DET3-F: 5'- TGGACCCCAACACAGATTGG-3' and -R: 5'-TGGATCAGGACGTCGAAAGC-3'), treated with exoSAPIT (Life Technologies) and analyzed by agarose gel electrophoresis. From 96 colonies, approximately 19 appeared to have a single-band deletion of the predicted size. Aliquots of the PCR reactions were sequenced with the F primer to confirm. To confirm homozygosity, PCR products from four candidates were cloned into a plasmid (StrataClone Blunt PCR Cloning Kit, Agilent) and individual colonies sequenced to identify each allele. Only one candidate, A7, exhibited the designed homozygous 76 bp deletion. A second colony with no deletion, E11, was selected to serve as unedited control.

### Bulk RNA sequencing analysis

Fastq files were downloaded, filtered with Fastp[6], aligned to human reference genome (hg38) with HISAT2[7], SNPs in mapped *KCNJ6* sequence were extracted using samtools mpileup[8] and bcftools to filter for quality. Ensembl VEP[9] was used to annotate SNPs.

### Single Cell RNA Sequencing

Dissociated cells were centrifuged at 200 x g for 5 minutes at 4°C. The supernatant was removed, and cells were washed twice with 1X HBSS (Thermo Fisher). After washes, cells were resuspended in 1% BSA (Bovine Serum Albumen, ThermoFisher)/1xPBS (ThermoFisher). Cells were counted and resuspended at 1200 cells/µl in 1% BSA/ 1xPBS and placed on ice. Single-cell cDNA libraries were generated using the Chromium Next GEM Single Cell 3' GEM Reagent Kits v3.1 (10X Genomics). Approximately 20,000 dissociated cells were loaded onto a cassette in the Chromium Controller with accompanying reagents to generate Gel Beads in Emulsions (GEMs). GEMs were subjected to reverse transcription to generate single-cell cDNA libraries before the oil emulsion was disrupted and cDNA purified using Dynabeads MyOne Silane (Thermo Fisher). cDNA was amplified by PCR for 11 cycles and purified using SPRIselect reagent (Beckman Coulter). cDNA fragmentation, A-tailing, and end repair was performed followed by adapter ligation for paired-end sequencing. An additional 11 cycles of PCR were performed to incorporate the sample index sequences and to amplify the libraries before samples were purified using SPRIselect reagent and used for sequencing. Libraries were shipped to Psomagen, Inc., for sequencing service. Fastq files were aligned with reference genome (mixed GRCh38and mm10) using CellRanger (10XGenomics) and then imported into a Seurat (v. 4.0.6) object[10]. A summary of human (GRCh38) and mouse (mm10) characteristics is shown in Supplemental Figure 2A and CellRanger parameters are provided in Supplemental Table 2. A cluster consistent with the co-cultured mouse glia was identified (Supplemental Figure 2B) and removed. Human cells were isolated for further analysis. Cells from individual subjects were identified using genome-wide SNPs that were expressed in the mRNA. Demuxlet[11] assigned "best guess" identification by cell barcode, which was imported as metadata into the human Seurat object. The number of identified cells was plotted for each cell line and treatment (Supplemental Figures 2C). Seurat clusters consistent with neuronal identity (Figure 1C, Supplemental Figure 2E) were selected for further analysis. Total numbers of neurons by subject and treatment are plotted in Supplemental Figure 2D. A minimum of 10 cells per subject and treatment were required for analysis—only subject 351 in the control condition fell below this threshold.

Differential gene expression was performed on gene counts aggregated by subject ("pseudobulk" analysis). To exclude sporadically-expressed genes, detectable expression was required in at least 25% of the cells in one group. After filtering, gene counts were summed for each cell line and condition. Tables of summed gene counts were analyzed by DESeq2[12] using the likelihood ratio test (LRT), comparing a model of groups to a reduced model. Contrasts were calculated and significance required an adjusted p-value of 0.05 or less and at least a 1.5-fold difference. Genes either up- or down-regulated were assessed for enriched gene ontology biological process terms using the enrichGO function from the clusterProfiler package[13]. Summary data are listed in Supplemental Figure 2.

**Fluorescent in situ hybridization**

iN cultures were fixed in 4% paraformaldehyde for 20 min at room temperature, followed by a series of ethanol dehydration/rehydration steps, permeabilization (with PBS-0.01% Tween buffer), blocking and overnight incubation with primary antibody against MAP2 (Millipore AB5543, 1:100 dilution). Prior to probe hybridization, samples were treated with hydrogen peroxide for 10 min and then with protease III (1:10) for 10 min. Human-specific *KCNJ6* probe was hybridized for 2 h, followed by standard amplification and color reaction with 520 Opal dye (Akoya Biosciences). MAP2 antibody was labeled with secondary antibody (Alexa 633), nuclei were stained with DAPI, and coverslips were mounted on slides using mounting medium (Thermo Fisher). Images were acquired on a Zeiss LSM 800 confocal microscope using a 40x objective. *KCNJ6* was quantified by manually counting fluorescent puncta within the MAP2/DAPI positive regions of individual neurons (defined as "somatic") or as puncta scattered along the MAP2-associated processes ("non-somatic"). Levels of *KCNJ6* were binned according to semi-quantitative histological scoring methodology by levels using Advanced Cell Diagnostics scoring criteria, where low expression of RNA target was defined to be 1-3 puncta per cell, intermediate 4-9, high 10-15 and very high >15 per cell.

**Electrophysiology**

Recordings were done in HEPES buffer consisting of (in mM): 140 NaCl, 5 KCl, 2 $CaCl_2$, 2 $MgCl_2$, 10 HEPES, 10 Glucose. Osmolality of the buffer was between 290-305 mOsm and the pH was adjusted to 7.4. Recordings were performed using borosilicate glass electrodes with resistances of 4-8 MΩ and intracellular solution containing the following (in mM): 126 K-Gluconate, 4 KCl, 10 HEPES, 4 ATP-Mg, 0.3 GTP-$Na_2$, 10 Phosphocreatine, with pH adjusted to 7.2 and osmolarity to 270−290 mOsm. After patching, neurons were allowed to rest for 5-7 min, or until the RMP drift stabilized. For GIRK2 channel recordings, an extracellular solution with high $K^+$ consisted of: 20 mM KCl, 140 mM NaCl, 2 mM $CaCl_2$, 2 mM $MgCl_2$ and 10 mM HEPES (pH 7.4). GIRK-mediated currents were measured at -40 mV membrane potential holding. Excitability of the neurons was measured by applying ramp (+200 pA) and step (-50 pA – 100 pA) depolarizing pulses while holding neurons at -65mV (current clamp mode). All recordings were performed at room temperature. Cells were excluded from analysis if access resistance changed by more than 20% during recording. For current injection studies, neurons were excluded if RMP was close to -63mV and the required current injection was less than 1 pA (absolute value). Any induced neurons whose access resistance was greater than 35 MΩ were also excluded.

**Calcium imaging**

Coverslips were mounted on a diamond-shaped chamber and placed on a Nikon Eclipse TE2000-U microscope with a 10X objective. GCaMP6f was excited using a 480 nm (Mic-LED-480A, Prizmatix), a HQ480/40x excitation filter, a Q505LP dichroic mirror, and a HQ535/50m emission filter (Semrock). Fluorescence was projected onto a sCMOS Zyla 5.5 camera (Andor) and sampled at a rate of 4.7 fps with a frame exposure of 200 ms at 160x120 pixels (4x4 binning). Light source and sCMOS camera were controlled with the Nikon Elements software (NIS-Elements AR 5.20.01).

Neurons were continuously perfused during fluorescence recording with ACSF with the following composition (in mM): NaCl 125, KCl 5, D-Glucose 10, HEPES-Na 10, $CaCl_2$ 3.1 and $MgCl_2$ 1.3. The pH was adjusted to 7.4 with HCl and osmolarity corrected with sucrose to 290-300 mOsm. Depolarizing solutions were prepared in ACSF with 10 or 50 µM glutamate, or with 18 mM KCl. Perfusion was gravity fed (flow rate of 0.065 ml/s). Solution exchanges were controlled with a ValveBank8 II (AutoMate Scientific Inc.). Calcium transients were recorded using a protocol based on glutamate and KCl perfusion, that consisted of 3 min baseline activity (ACSF), 3 x 30 s pulses of 10 µM glutamate (30 s ACSF between pulses), one 30 s pulse of 50 µM glutamate, 2 min of ACSF wash, 3 x 30 s pulses of 18 mM KCl (30 s ACSF between pulses), and a final ACSF wash for 5 min (Fig. 4C).

ROI segmentation of GCaMP6f-expressing neurons, raw fluorescence extraction and background correction were performed with Nikon Elements software. ΔF/F was calculated with the formula $(F_t - F_{min})/F_{min}$, where $F_t$ is the raw fluorescence at time t and $F_{min}$ is the minimum fluorescence for the entire trace. The resulting trace was then denoised with a low-pass Butterworth filter and baseline corrected for drift using an adaptative iteratively reweighted Penalized Least Squares (AirPLS) based algorithm [14], in RStudio (R version 4.0.3). Peak detection was performed using a custom R script that considered neuron action potential-derived $Ca^{2+}$ spikes that had the following criteria (for a framerate acquisition of 4.7 fps): duration < 30 frames, rise phase >= 2 frames, fall phase >= 5 frames, rise phase < fall phase, peak height > 5*SD, and peak height > 5*max background signal. The analysis pipeline is depicted in Supp. Fig. 6. Statistical analysis was performed in GraphPad Prism 8.4.3 or R version 4.0.3.

**Image analysis**

Regions of Interest (ROIs) were created for each cell by encapsulating the cell body using the cell magic wand tool. The cell body was then quantified for area and shape. The NeurphologyJ Fiji macro[15] was utilized for detection and quantification of extended neuronal morphometry. A threshold was selected manually so that most neurites are well-defined without significant particle detection or noise. GIRK2 puncta quantification used trainable Weka Segmentation[16] to segment the image to establish two classes. One class identified pixels that corresponded to GIRK2 puncta for intensity, circularity, and size. A second class identified pixels that did not correspond to puncta, such as noise from neurons and background. The classifier was trained multiple times until there was a high confidence that the pixels segmented into class one had a high (90%) confidence interval denoted by probability maps. The segmented image was adjusted to remove any false negatives. The class one binary image was then used for puncta cytometry.

### References


1. Weissbein U, Schachter M, Egli D, Benvenisty N. Analysis of chromosomal aberrations and recombination by allelic bias in RNA-Seq. *Nature communications* 2016; **7:** 12144.
2. Yang N, Chanda S, Marro S, Ng Y-H, Janas JA, Haag D *et al.* Generation of pure GABAergic neurons by transcription factor programming. *Nature Methods* 2017; **14**(6)**:** 621-628.
3. Kreitzer FR, Salomonis N, Sheehan A, Huang M, Park JS, Spindler MJ *et al.* A robust method to derive functional neural crest cells from human pluripotent stem cells. *Am J Stem Cells* 2013; **2**(2)**:** 119-131.
4. Miyaoka Y, Chan AH, Judge LM, Yoo J, Huang M, Nguyen TD *et al.* Isolation of single-base genome-edited human iPS cells without antibiotic selection. *Nature methods* 2014; **11**(3)**:** 291-293.
5. Ran FA, Hsu PD, Wright J, Agarwala V, Scott DA, Zhang F. Genome engineering using the CRISPR-Cas9 system. *Nature Protocols* 2013; **8**(11)**:** 2281-2308.
6. Chen S, Zhou Y, Chen Y, Gu J. fastp: an ultra-fast all-in-one FASTQ preprocessor. *Bioinformatics (Oxford, England)* 2018; **34**(17)**:** i884-i890.
7. Kim D, Paggi JM, Park C, Bennett C, Salzberg SL. Graph-based genome alignment and genotyping with HISAT2 and HISAT-genotype. *Nature Biotechnology* 2019; **37**(8)**:** 907-915.
8. Danecek P, Bonfield JK, Liddle J, Marshall J, Ohan V, Pollard MO *et al.* Twelve years of SAMtools and BCFtools. *GigaScience* 2021; **10**(2).
9. McLaren W, Gil L, Hunt SE, Riat HS, Ritchie GRS, Thormann A *et al.* The Ensembl Variant Effect Predictor. *Genome Biology* 2016; **17**(1)**:** 122.
10. Butler A, Hoffman P, Smibert P, Papalexi E, Satija R. Integrating single-cell transcriptomic data across different conditions, technologies, and species. *Nat Biotechnol* 2018; **36**(5)**:** 411-420.
11. Kang HM, Subramaniam M, Targ S, Nguyen M, Maliskova L, McCarthy E *et al.* Multiplexed droplet single-cell RNA-sequencing using natural genetic variation. *Nat Biotechnol* 2018; **36**(1)**:** 89-94.
12. Love MI, Huber W, Anders S. Moderated estimation of fold change and dispersion for RNA-seq data with DESeq2. *Genome Biol* 2014; **15**(12)**:** 550.
13. Wu T, Hu E, Xu S, Chen M, Guo P, Dai Z *et al.* clusterProfiler 4.0: A universal enrichment tool for interpreting omics data. *Innovation (N Y)* 2021; **2**(3)**:** 100141.
14. Zhang ZM, Chen S, Liang YZ. Baseline correction using adaptive iteratively reweighted penalized least squares. *Analyst* 2010; **135**(5)**:** 1138-1146.
15. Ho S-Y, Chao C-Y, Huang H-L, Chiu T-W, Charoenkwan P, Hwang E. NeurphologyJ: An automatic neuronal morphology quantification method and its application in pharmacological discovery. *BMC Bioinformatics* 2011; **12**(1)**:** 230.
16. Arganda-Carreras I, Kaynig V, Rueden C, Eliceiri KW, Schindelin J, Cardona A *et al.* Trainable Weka Segmentation: a machine learning tool for microscopy pixel classification. *Bioinformatics* 2017; **33**(15)**:** 2424-2426.

**Supplemental Figure 1**

A. **Confirmation of pluripotency in iPSC lines**

Immunocytochemical detection of pluripotency markers Oct4 and TRA-1-60 in four of the eight iPSC lines (Table 1). All iPSC lines exhibited these markers and the characteristic colony morphology.

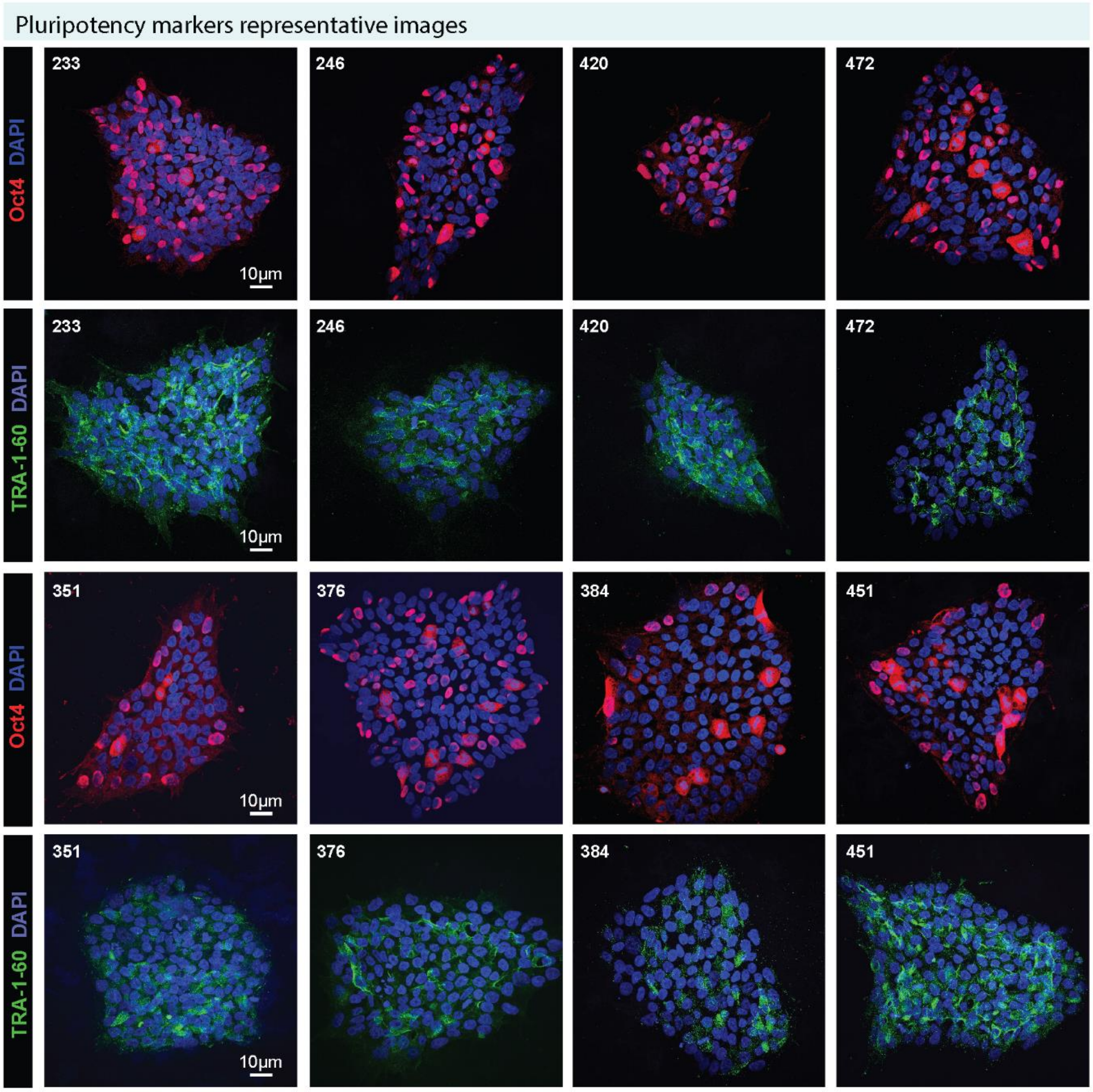

### B. eKaryotyping

Bulk RNAseq data from all eight iPSC lines were processed as described for e-Karyotyping analysis (60). Top, uniformity of read detection across the genome. Bottom, allelic ratio plots. No anomalies were detected.

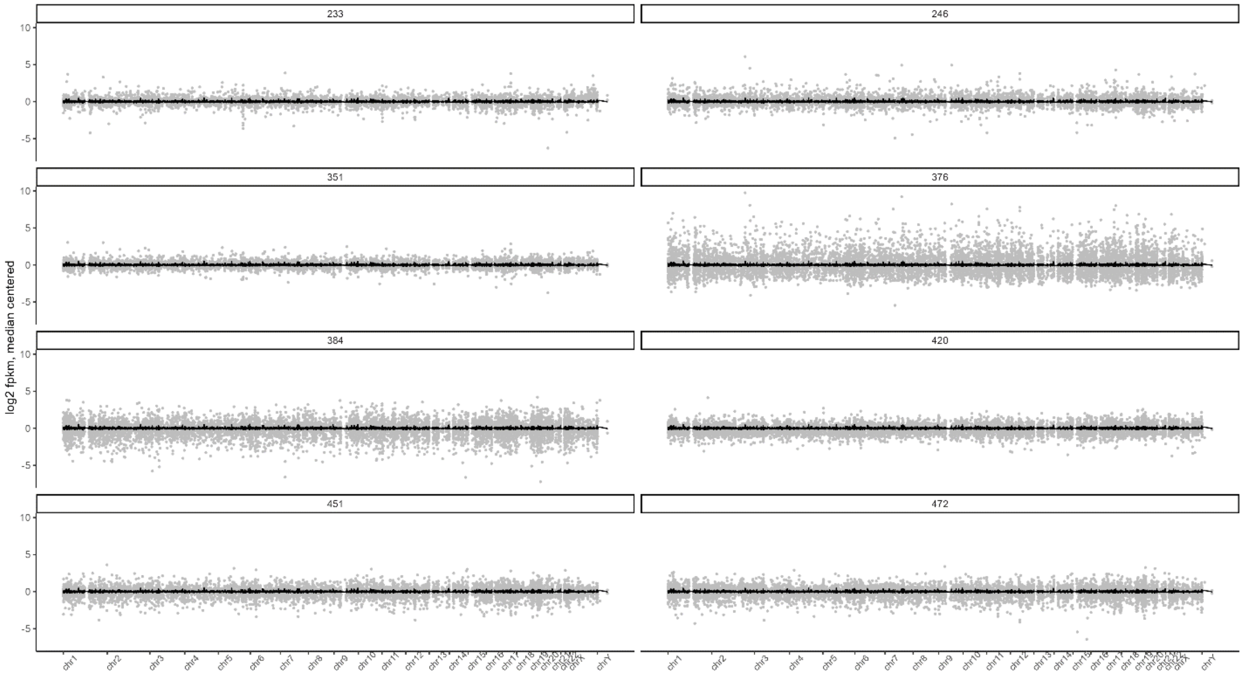

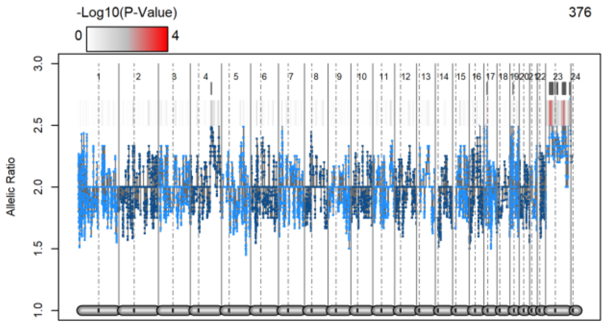
-Log10(P-Value)
0 4
376
Allelic Ratio
3.0 2.5 2.0 1.5 1.0
1 2 3 4 5 6 7 8 9 10 11 12 13 14 15 16 17 18 19 20 21 22 23 24

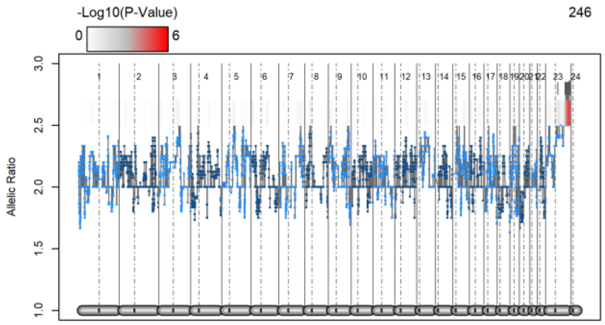
-Log10(P-Value)
0 6
246
Allelic Ratio
3.0 2.5 2.0 1.5 1.0
1 2 3 4 5 6 7 8 9 10 11 12 13 14 15 16 17 18 19 20 21 22 23 24

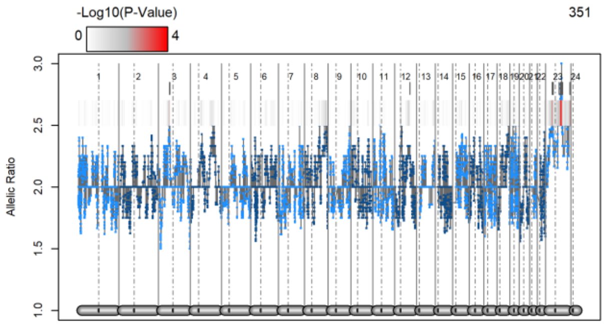
-Log10(P-Value)
0 4
351
Allelic Ratio
3.0 2.5 2.0 1.5 1.0
1 2 3 4 5 6 7 8 9 10 11 12 13 14 15 16 17 18 19 20 21 22 23 24

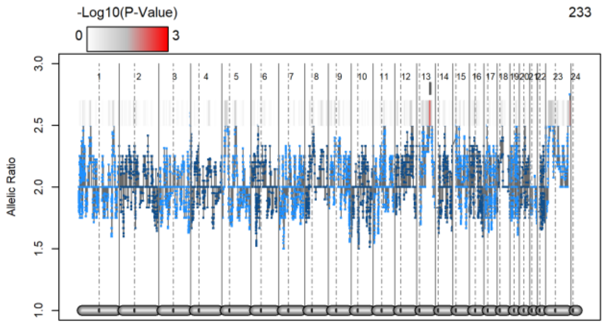
-Log10(P-Value)
0 3
233
Allelic Ratio
3.0 2.5 2.0 1.5 1.0
1 2 3 4 5 6 7 8 9 10 11 12 13 14 15 16 17 18 19 20 21 22 23 24

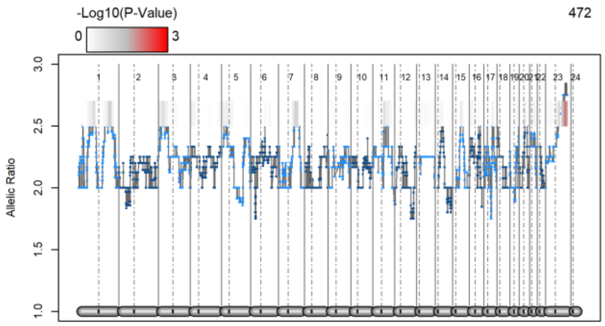
-Log10(P-Value)
0 3
472
Allelic Ratio
3.0 2.5 2.0 1.5 1.0
1 2 3 4 5 6 7 8 9 10 11 12 13 14 15 16 17 18 19 20 21 22 23 24

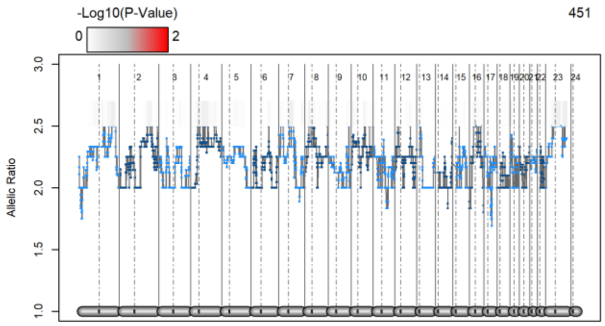
-Log10(P-Value)
0 2
451
Allelic Ratio
3.0 2.5 2.0 1.5 1.0
1 2 3 4 5 6 7 8 9 10 11 12 13 14 15 16 17 18 19 20 21 22 23 24

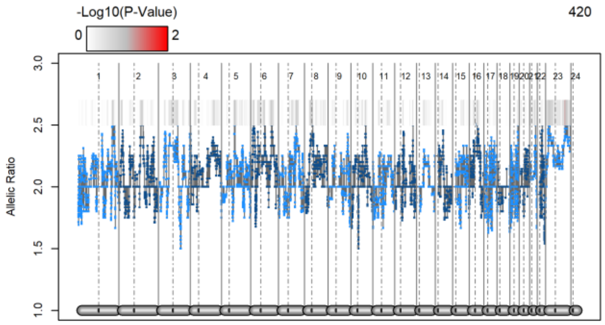
-Log10(P-Value)
0 2
420
Allelic Ratio
3.0 2.5 2.0 1.5 1.0
1 2 3 4 5 6 7 8 9 10 11 12 13 14 15 16 17 18 19 20 21 22 23 24

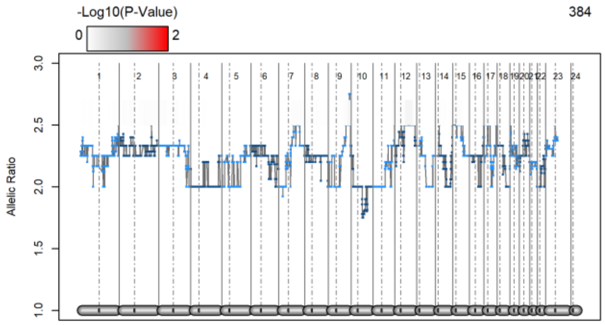
-Log10(P-Value)
0 2
384
Allelic Ratio
3.0 2.5 2.0 1.5 1.0
1 2 3 4 5 6 7 8 9 10 11 12 13 14 15 16 17 18 19 20 21 22 23 24

C. **Correlation of SNPs in KCNJ6 3'UTR.** Using 5 EUR populations we assessed linkage disequilibrium pattern (LDLINK-NCI; https://ldlink.nci.nih.gov/?tab=ldassoc) of SNPs within the range identified in our analyses. In red is plotted the $R^2$ (ranging from 0.882 to 0.996) and blue is the D' (ranging from 0.75 to 0.996). Results confirm a strong linkage disequilibrium in the region.

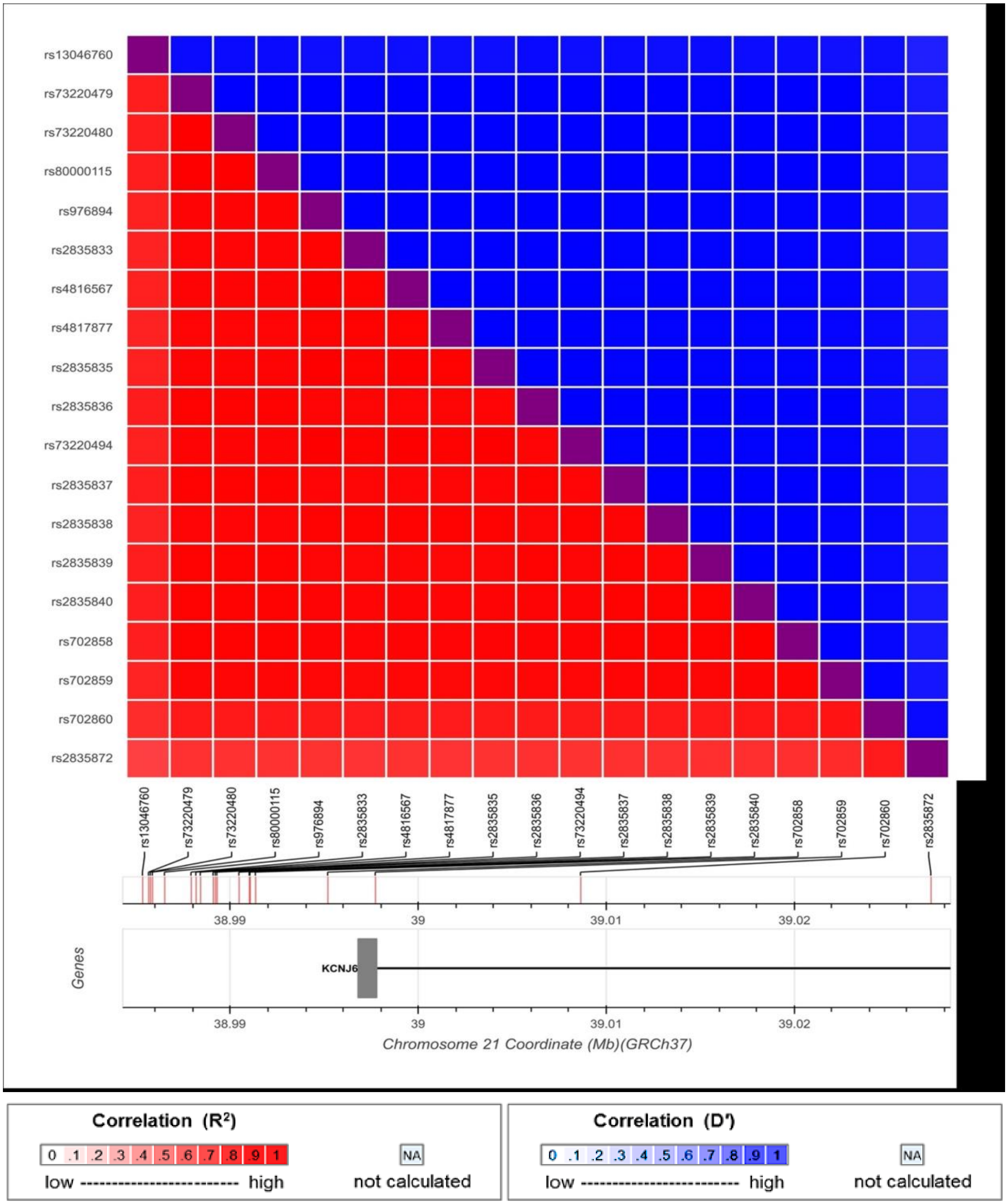

**Supplemental Figure 2**

A. **scRNAseq summary table.** For each sample, the numbers of genes aligning with human (GRCh38) and mouse (mm10) reference genomes are shown. First character: C = Unaffected group, A = Affected group. Second character: C = Control and 7 = 7d of IEE.

| Sample | Total Reads | GRCh38 Median Genes per Cell | mm10 Median Genes per Cell | GRCh38 Total Genes Detected | mm10 Total Genes Detected | GRCh38 Median UMI Counts per Cell | mm10 Median UMI Counts per Cell |
|---|---|---|---|---|---|---|---|
| CC | 459,677,576 | 2,997 | 1,288 | 30,109 | 15,224 | 7,961 | 3,726 |
| C7 | 347,985,342 | 1,407 | 2,395 | 30,107 | 19,138 | 2,422 | 6,979 |
| AC | 377,021,443 | 1,735 | 514 | 29,740 | 14,478 | 3,641 | 1,199 |
| A7 | 361,613,420 | 2,240 | 3,009 | 29,489 | 15,431 | 4,668 | 9,412 |

B. **Identification of human and mouse cells.** A tSNE plot of all cells shows the clusters with predominantly mouse cells (cyan) distinct from those with predominantly human cells (red). Mouse cells were excluded from all subsequent analyses and tSNE plots were re-calculate on only human gene expression.

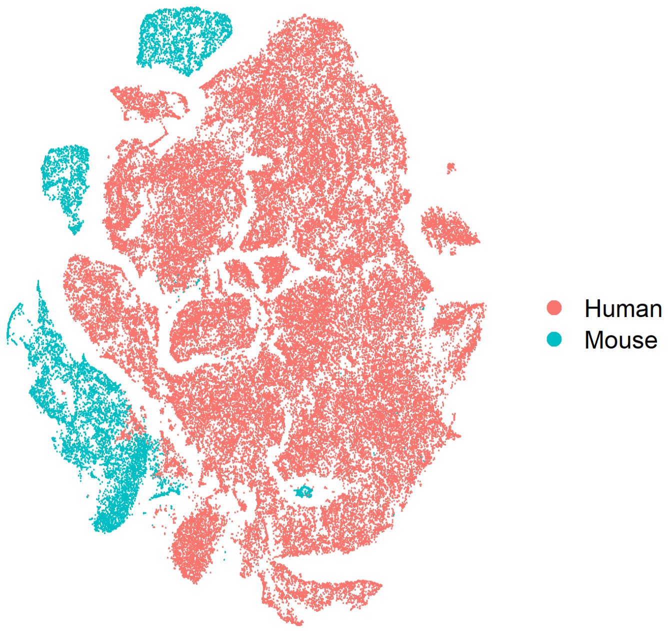

C. **scRNAseq cell distribution by subject and treatment.** Human cells were assigned to individual subject identifiers using SNP data from GWAS studies. Plotted are the numbers of cells per subject, either from the untreated control samples or from the samples exposed to 7 d of IEE ("Alcohol").

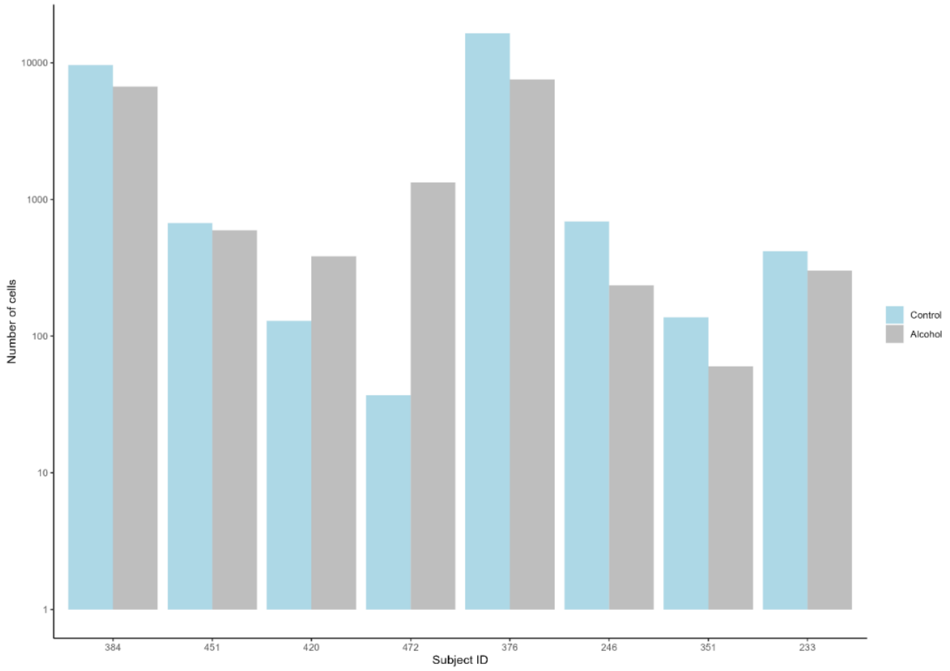


D. **Numbers of selected neurons by subject and treatment.** After identifying the neuron cluster (Fig. 1C), the numbers of neurons per subject and treatment group is plotted. The dashed line shows the acceptable minimum threshold to be included for pseudo-bulk analysis.

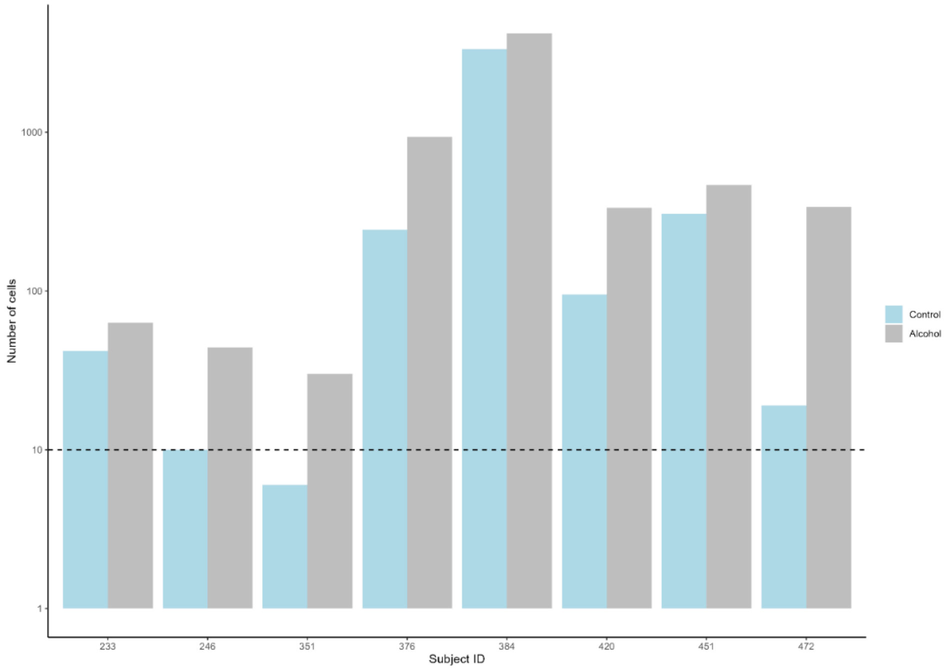

E. **Genes enriched in induced neurons.** As in Fig. 1C, for each gene, colored dots in the tSNE plot indicates expression levels for each cell. Neuron structural proteins (MAPT, MAP2, TUBB3) are expressed widely but are especially strong in the neuron cluster (lower left, see key in Fig. 1C). Several members of the glutamate receptor families are similarly enriched (NMDA: GRIN2A, GRIN2B, GRIA2, GRIA4) as is one example of the kainate family (GRIK2).

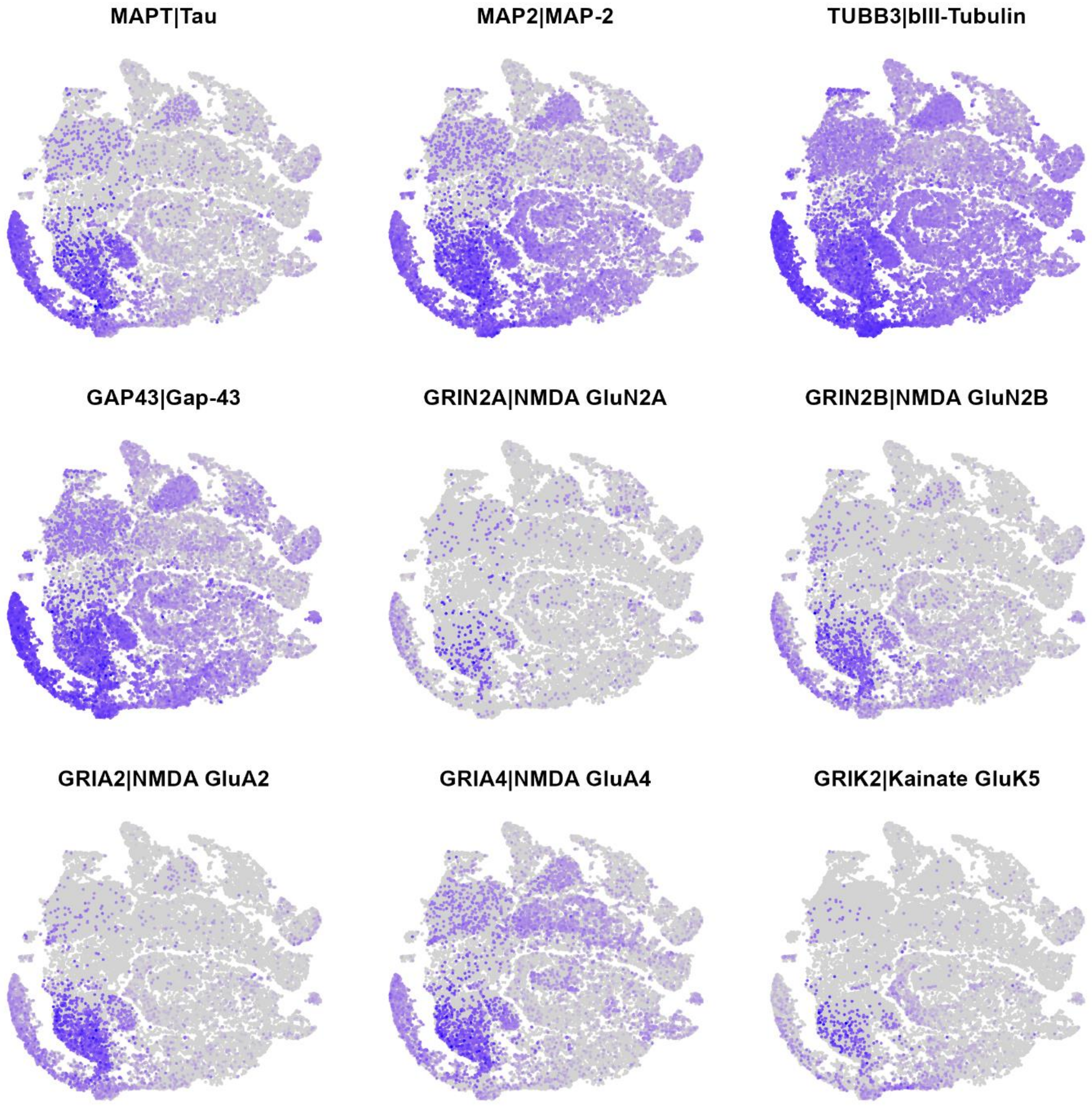

**Supplemental Figure 3**

**A. Biological process (BP) gene ontology terms enriched in up-regulated genes (AF > UN), grouped as a treeplot.**

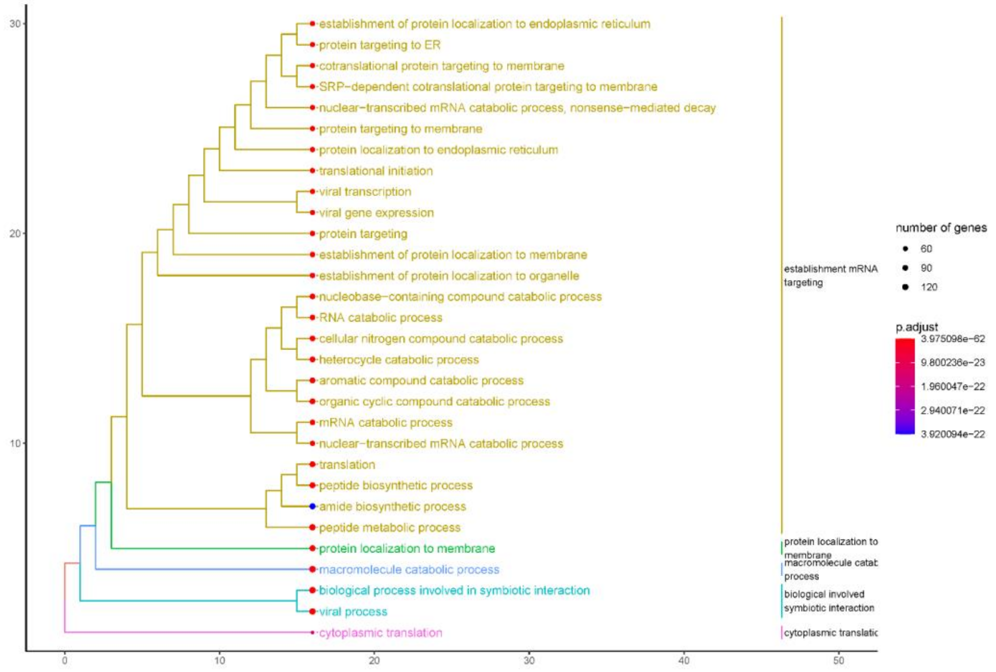


**B. Biological process gene ontology terms enriched in down-regulated genes (AF < UN), grouped as a treeplot.**

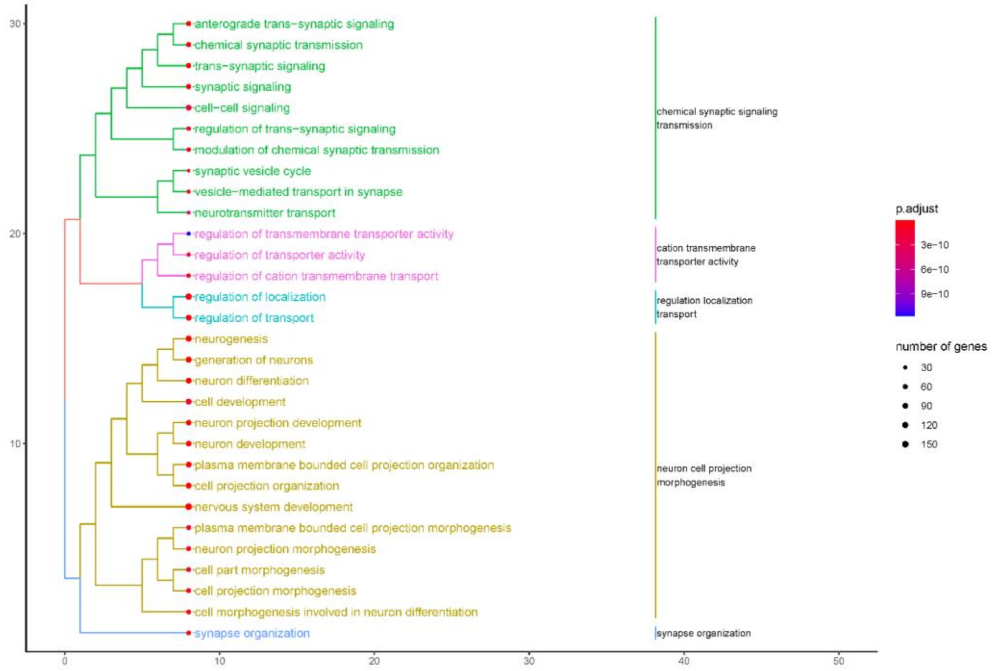

**Supplemental Figure 4**

**GIRK2 immunocytochemical detection in primary mouse neuron cultures and antibody specificity demonstrated by silencing or knockout.** (A) Alternate GIRK2 expression patterns observed in mouse neurons. Human iN (Fig. 2B) more closely match a pattern with expression primarily in neural processes (Fig. 2A). A portion of mouse neurons label more uniformly throughout the cell, as shown here. GIRK2 puncta do not completely colocalize with SYN1. (B) Human iN cultures infected with lentiviral-encoded shRNAs targeting KCNJ6 exhibit reduced GIRK2 immunoreactivity. (C) A human iPSC line CRISPR/Cas9 edited to introduce frameshift mutations in both alleles of KCNJ6 exhibit a lack of GIRK2 immunoreactivity. As control, lentiviral overexpression of KCNJ6 restores immunoreactivity.

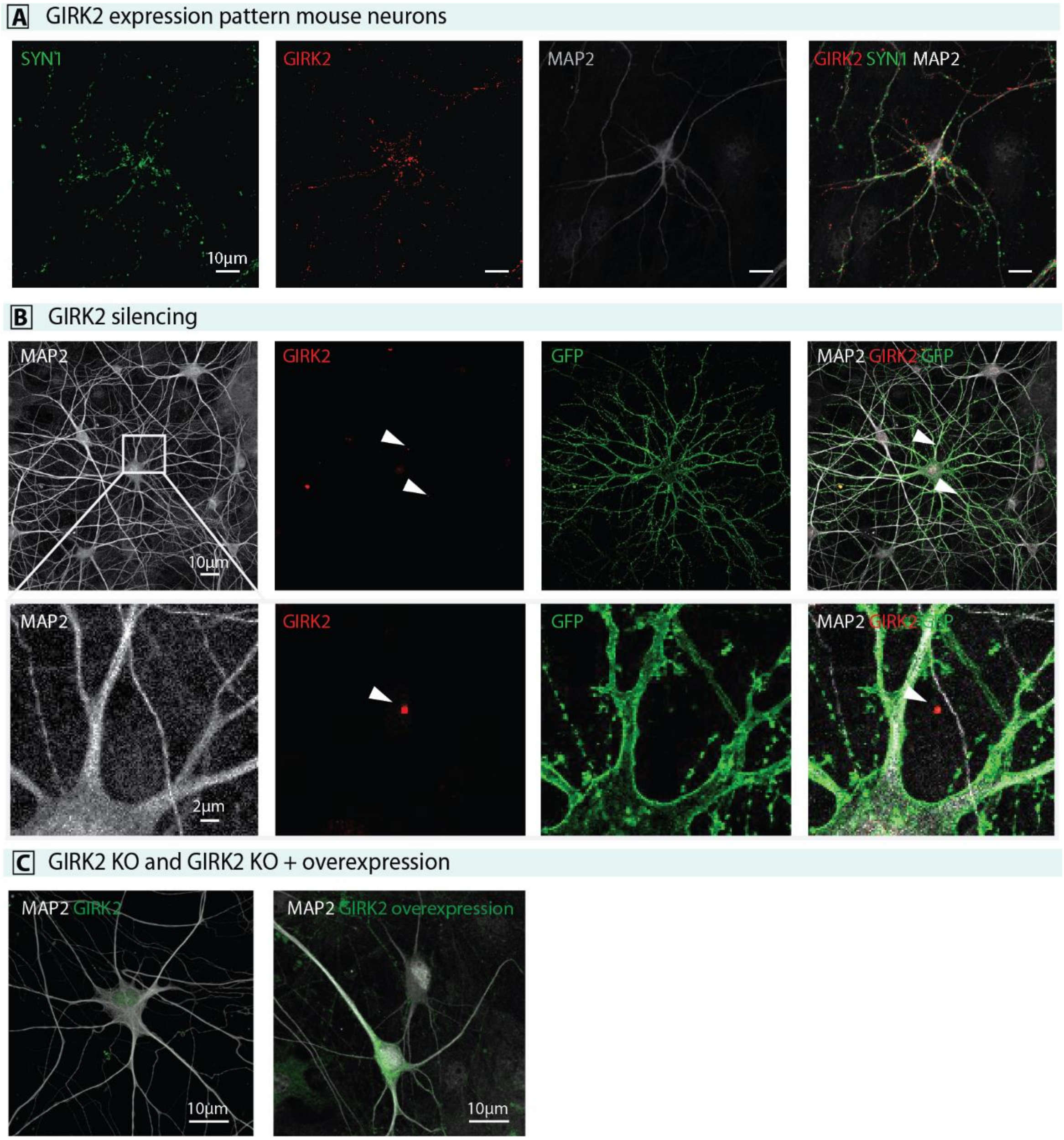

**Supplemental Figure 5**

A. GIRK2 expression co-localizes with βIII-tubulin (Fig. 2) and MAP2, but does not directly colocalize with VGLUT2, a marker of synaptic vesicles in glutamatergic neurons.

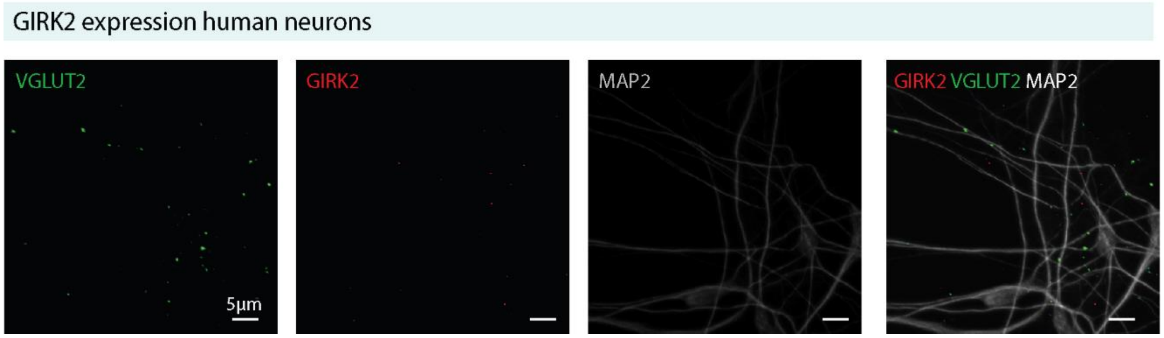


B. GIRK2 does not colocalize with synaptic marker Syn1. This enhanced image was produced using Airyscanning to improve GIRK2 resolution.

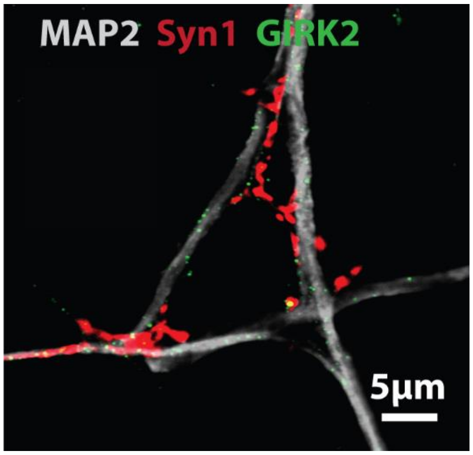

**Supplemental Figure 6. Calcium imaging analysis pipeline and evaluation of neuronal excitability in iN populations by individual subject.**

**A**. Analysis pipeline and representative calcium imaging raw fluorescence and dFoF traces of affected individuals 233 and 246, and unaffected individuals 420 and 472. **a, b.** The ROIs are selected from the GCaMP6f-expressing neurons in the imaging field; **c, d, e, f.** The raw fluorescence data of each ROI is extracted, noise-filtered and baseline corrected using an AirPLS algorithm to generate the dFoF traces. A peak-detection algorithm is used to extract the peaks from the dFoF, corresponding to calcium spikes. The vertical dotted lines indicate the calcium spikes detected after analysis. **B, C, D.** Number of spikes per ROI per minute fired by neurons from each line of unaffected (grey) and affected individuals (blue) during baseline (**B**, spontaneous activity), glutamate pulse 1 (**C**, glutamate-elicited activity) and KCl pulse 1 (**D**, KCl-elicited activity). The sample size is depicted in the bar of each line as number of Nng2-neuron batches/number of experiments/number of neurons. Differences between lines were evaluated by one-way ANOVA (Brown-Forsythe test for non-equal SD) and Tukey's multiple comparisons test (**p < 0.01, ****p < 0.0001).

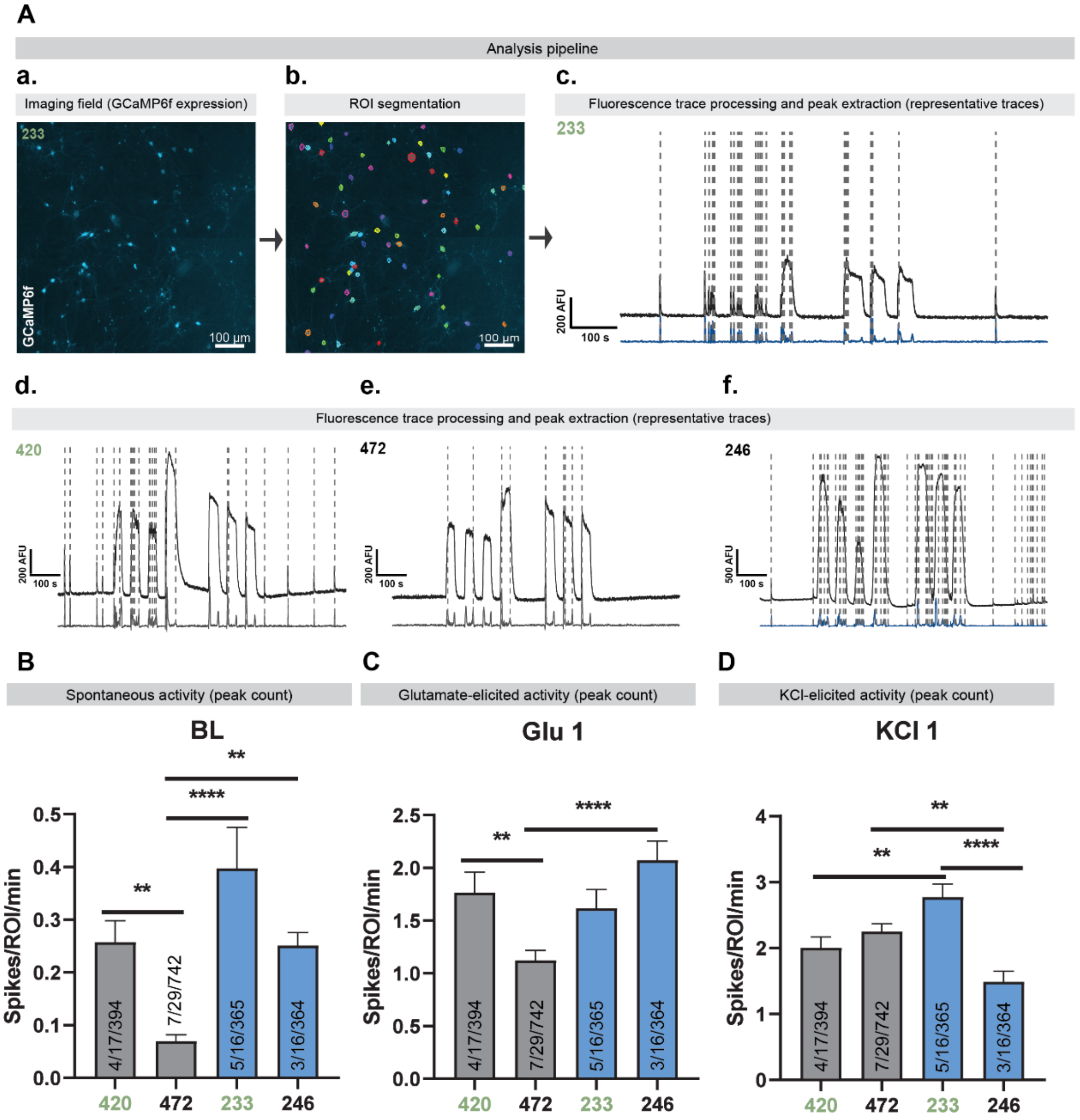

**Supplemental Figure 7**

**A. Ethanol decay over 7 days of IEE.** Sampling medium from two lines (351 and 384), IEE produced a mean concentration of 15.4 mM ± 1.2 (SEM) over 7 days, with a half-life of 14.46 h (95% CI: 13.45 to 15.62 h).

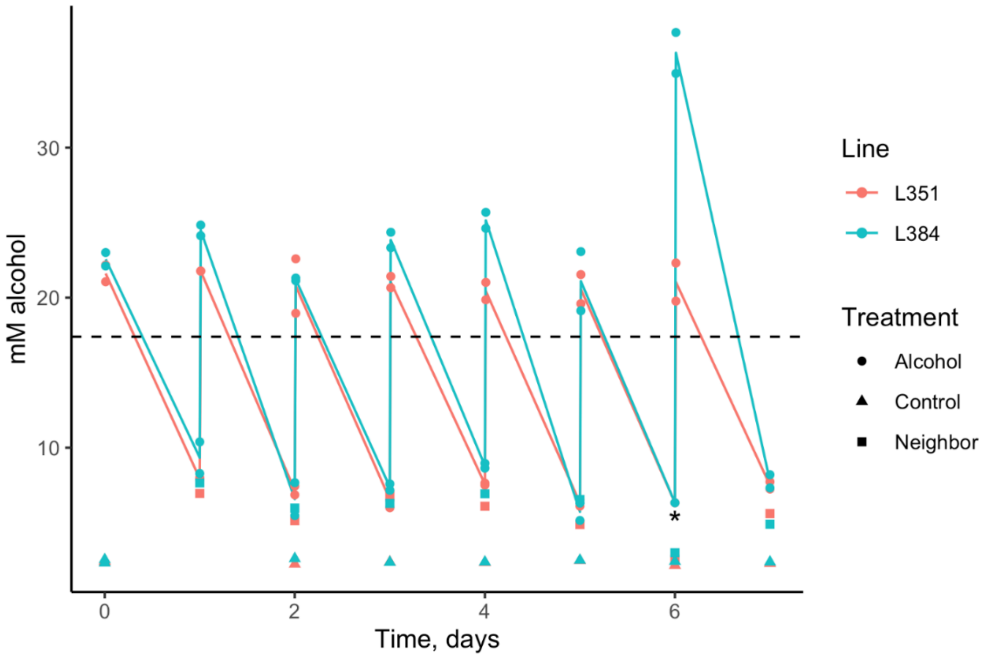


**B. Regression analysis of decay from time of medium supplementation.**

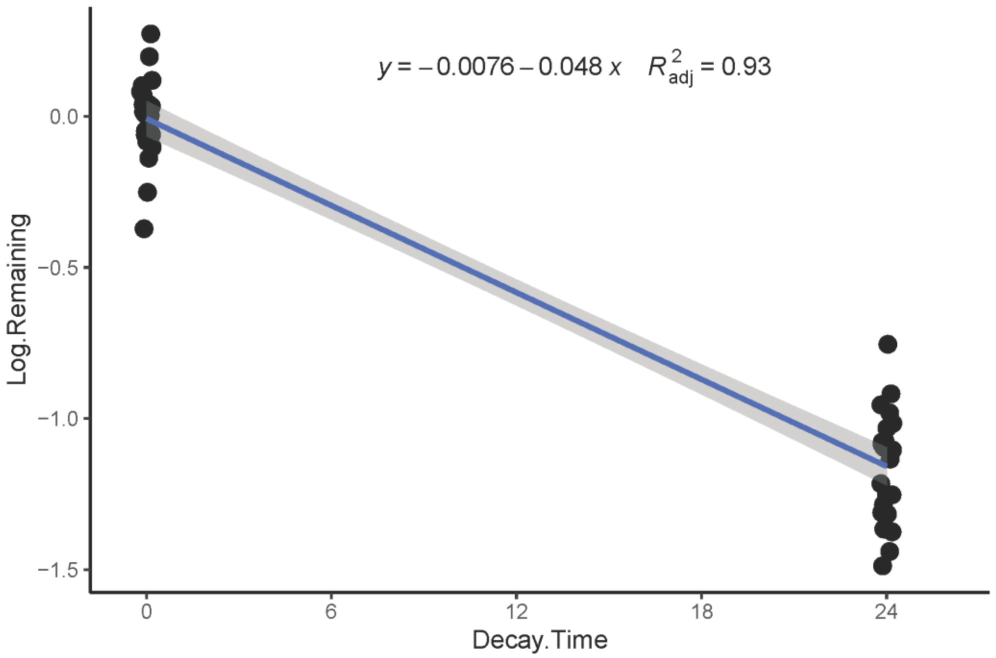
